# Dynamics in microbial communities associated with the development of soil fatigue in banana

**DOI:** 10.1101/2025.05.14.653785

**Authors:** David-Dan Cohen, Adi Faigenboim, Idan Elingold, Yonatan Sher, Navot Galpaz, Dror Minz

**Affiliations:** Department of Soil Chemistry, Plant Nutrition and Microbiology, Institute of Soil, Water and Environmental Sciences, Agricultural Research Organization, Volcani Center, Beit Dagan, Israel; Department of Agroecology and Plant Health, Robert H. Smith Faculty of Agriculture, Food and Environment, Hebrew University of Jerusalem, Jerusalem, Israel; Institute of Plant Science, Agricultural Research Organization, Volcani Center, Beit Dagan, Israel; Jordan Valley Banana Research Station, Zemach 15132, Israel; Northern Research and Development, MIGAL Galilee Research Institute, Kiryat Shmona, Israel

**Keywords:** Banana plants, 16S, ITS, Bioinformatic, Soil fatigue, Pathogen, Beneficial microorganisms, Microbial ecology, Soil chemistry, Microbiome

## Abstract

Soil fatigue, well-documented in various crops, presents a significant challenge to banana production by causing fast and then gradual declines in plant growth and yield over years of cultivation. Despite its impact on profitability, the underlying mechanisms driving soil fatigue remain poorly understood; however, a strong link to shifts in the soil microbiome has been suggested. We investigated the dynamics of microbial communities in relation to soil fatigue, using a novel semi-controlled outdoor experimental system. Soils at different stages of fatigue (0 to 42 months of banana cultivation) were generated in large containers filled with initially healthy soil. Banana plants grown in these soils were replaced with new plants which showed soil age-dependent growth. Three months postplanting, soil and root samples were collected for analyses of soil parameters and microbial community composition using bacterial (16S) and fungal (ITS) amplicon sequencing. We identified minor age-related shifts in mainly pH, potassium, and organic matter in the soil. While alpha diversity remained unchanged, significant shifts in bacterial and fungal community composition were observed in fatigued soils. Notably, the relative abundance of bacterial families such as *Flavobacteriaceae, Pseudomonaceae*, and *Acidibacter* increased, as did some fungal taxa (many from groups with known pathogens)-*Ceratobasidiaceae* (including *Rhizoctonia)*, *Dothideomycetes*, and *Stachybotryaceae*. Simultaneously, the relative abundance of bacterial families with known beneficial members, including *Gemmatimonadaceae, Moraxellaceae, Sphingomonadaceae*, and *Azospirillaceae*, as well as symbiotic fungal taxa such as *Glomeraceae* and *Lasiosphaeriaceae*, declined. Thus, soil fatigue may be correlated to proliferation of pathogenic populations and a loss of beneficial microorganisms.

## 1. Introduction

The soil microbiome, comprising a diverse community of bacteria, fungi, viruses, and protists, is fundamental to plant health and productivity. Interactions between plants and their microbiomes shape key ecological and agricultural processes, including plant nutrition [1–4], disease suppression [5–7], and tolerance to abiotic stress [8–10]. Within the rhizosphere (the soil adjacent to, and affected by the plant roots; Hartmann et al., 2008), a complex web of interactions connects the roots, soil, and microbiomes [12, 13].

One of the significant challenges in modern agriculture is ‘soil fatigue’, which arises from repeated monoculture or continuous cultivation of the same crop on the same land [14, 15]. This phenomenon is closely linked to alterations in the soil and rhizosphere microbiomes [16–18] and the accumulation of pathogenic microorganisms in the soil [19–22]. Maintaining soil biodiversity is vital for preserving soil health [23] and achieving sustainable agricultural practices [24, 25]. Investigating the roles of soil and plant microbiomes in promoting plant health is key to developing strategies to mitigate the detrimental effects of soil fatigue [14, 26–28].

Research indicates that the decline in soil biodiversity caused by soil fatigue negatively impacts crop productivity and soil health [29–31]. High-throughput sequencing analyses have identified specific microbial taxa associated with soil fatigue and highlighted their potential roles in reduced plant growth [32, 33]. Long-term monitoring of soil age and microbial community dynamics [18, 34] provides critical insights into the mechanisms driving soil fatigue. This issue is particularly acute in banana cultivation, where soil fatigue significantly reduces productivity and increases production costs, posing a substantial threat to the global banana industry (Bonavita, 2022).

Most studies on soil fatigue in bananas and other crops compare microbial communities in plants grown at different times across geographically distinct fields [14, 26, 27] or under varied farming practices [18, 34]. However, such studies lack the ability to fully control experimental variables and to monitor, in real time, shifts in the microbiome within the same system.

In this study, we investigated the impact of soil age on banana growth and the associated development of soil and rhizosphere microbiomes. We designed an experimental system using identical soils of varying ages, coupled with banana plants at the same growth stage. These systems were sampled at predetermined intervals to enable controlled, real-time microbiome analyses. Our approach provides a novel perspective on understanding soil fatigue and its effects on plant growth and microbiome dynamics.

## 2. Materials and methods

### 2.1. Sampling site and experimental design

The study was carried out at the Tzemach experimental field, which is situated near the southern coast of the Sea of Galilee (<u>32.704545, 35.581159</u>). We filled 24 plastic containers of 1.5 m^3^ capacity with healthy soil (after 9 years of growing grapefruit in an orchard), which is considered optimal for banana cultivation (Fig. 1). Every 6 months from September 2015 to March 2019, three randomly selected containers were each planted with three banana plants [*Musa acuminata* (AAA group, cv. Grand Nain)]. These plants were maintained under standard agricultural practices, including fertigation using a fertilizer mix of N/P/K, supplemented with 6% (w/v) ammonium nitrate (NH_₄_NO_₃_), 1% (w/v) phosphoric acid (H_₃_PO_₄_), and 9% (w/v) potassium chloride (KCl). Unplanted containers were kept biologically active by sowing wheat annually in December. Irrigation and fertilization were uniformly applied to all containers, with each receiving 3.7 m³ of water annually, along with 67 g potassium, 101 g phosphorus, and 11 g nitrogen.

**Fig. 1.**
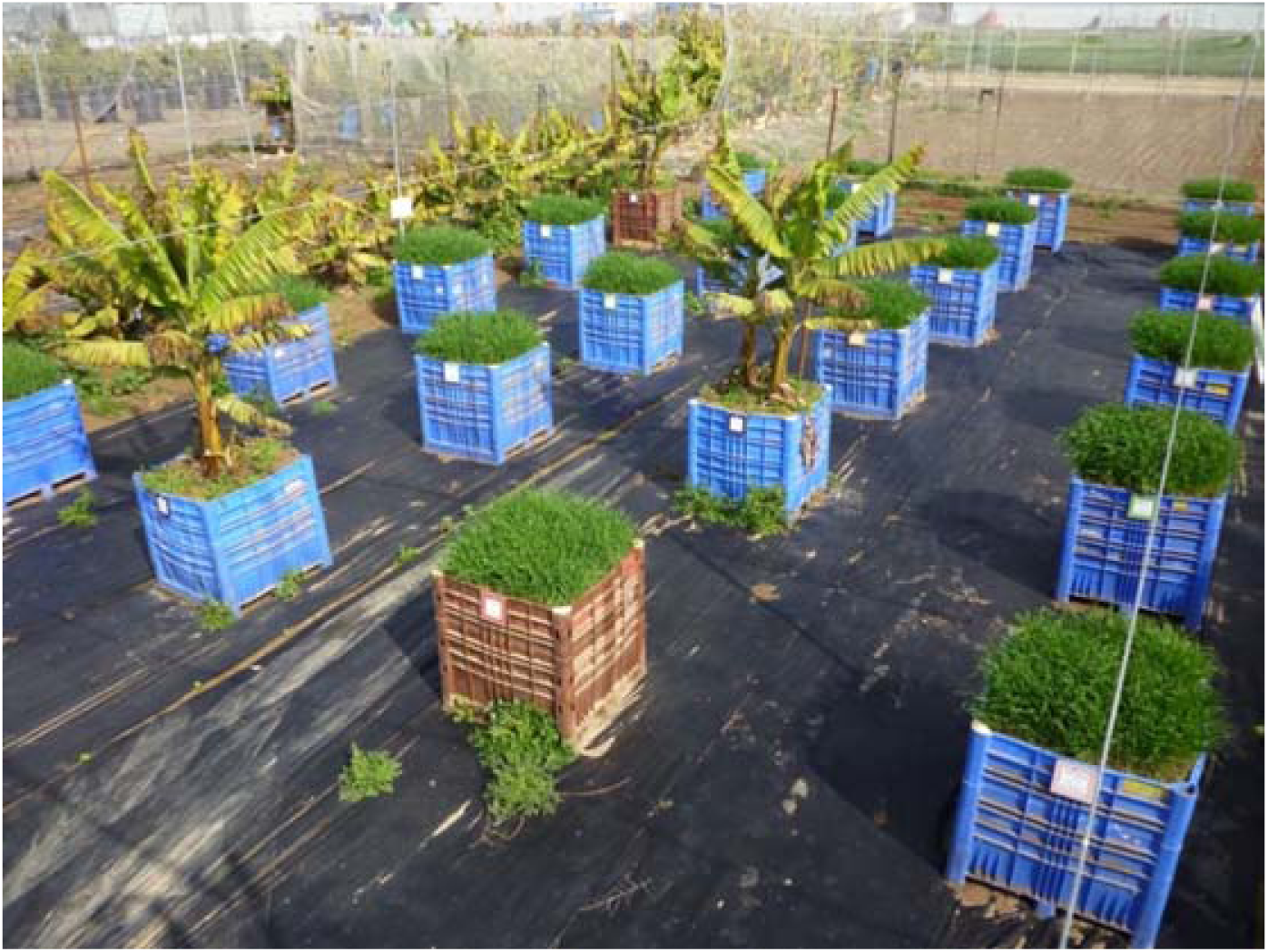
Experimental field. The experiment was composed of 24 containers, 1.5 m3 in size, that mimicked the soil fatigue phenomenon. Photo taken in February 2016, after only 3 containers out of the 24 were planted. Wheat was sown in the other containers in December 2015. These containers were later gradually planted with banana plants.

By March 2019, this gradual planting resulted in containers with banana plants (and their associated soils) of different ages: 42, 36, 30, 24, 18, 12, and 6 months, depending on the initial planting date (September 2015, March 2016, September 2016, March 2017, September 2017, March 2018, and September 2018, respectively). Three containers remained unplanted, maintaining a soil age of 0 months. In June 2019, all plants were removed. Then three new banana plants were planted in each container, ensuring that all plants were of identical age but growing in soils of different ages. After 3 months of growth, in September 2019, we measured plant height and the area of the third leaf-the latter considered representative for various measurements in banana studies. Plants were then removed using an excavator, and root, rhizosphere, and soil samples were collected from each container at a depth of 20 cm. Samples were immediately placed in liquid nitrogen and transported to the laboratory, where they were stored at −80 °C for future DNA isolation and analysis. In addition, the annual banana yield per hectare from commercial plantations in the Jordan Valley region was analyzed over a 6-year period. The average yield from these plots was calculated for each crop cycle.

### 2.2. Analysis of soil chemical parameters

In June 2019, immediately after the removal of the first set of banana plants and before planting the second set, soil samples were collected for chemical analyses according to the age of the soil. The aim of this evaluation was to determine whether changes in soil chemical parameters might affect the growth and development of the new banana plants. Sampling was conducted only for containers with soil aged 0, 6, 18, 30, and 42 months. The soil chemical parameters were measured at Zemach Agriculture Technologies Ltd. (Zemach, Israel), and included: pH, electrical conductivity (dS m^−1^), chlorine (mg L^−1^), sodium (mg L^−1^), calcium & magnesium (mEq L^−1^), available potassium (extracted with CaCl_₂_, mg L^−1^), available iron (mg kg^−1^), available zinc (mg kg^−1^), available manganese (mg kg^−1^), available boron (mg L^−1^), ammonium (mg kg^−1^), nitrate (mg kg^−1^), available phosphorus (Olsen P method, mg kg^−1^), sodium adsorption ratio (SAR), total organic carbon (TOC), and UV absorbance at 254 nm (as a measure of organic matter; Ahumada et al., 2001; Sirén et al., 1997). All measurements were carried out according to standard procedures. Multiple comparisons, using Student’s t-test, were performed for the outcomes of analyses of variance conducted with JMP 16 Pro software (SAS Institute Inc., Cary, NC, USA). All data are presented as means.

### 2.3. DNA extraction, PCR and sequencing

DNA was extracted from 300 mg soil using a FastPrep-24 bead beater (MP Biomedicals, Santa Ana, CA, USA) with Exgene soil DNA mini kit (GeneAll, Seoul, Korea) following the manufacturer’s instructions. DNA was eluted with 50 µL of elution buffer, and its concentration and purity were measured in a Qubit 2.0 fluorometer (Waltham, MA, USA). It was kept frozen at −20 °C until further analysis.

Amplicons were generated using a two-stage PCR-amplification protocol, as described previously [38]. All primers contained 5’ common sequence tags (CS1 and CS2). Fungal internal transcribed spacer (ITS) regions were amplified with primers ITS1F and ITS2R (CS1_ITS1F: ACACTGACGACATGGTTCTACACTTGGTCATTTAGAGGAAGTAA; CS2_ITS2R: TACGGTAGCAGAGACTTGGTCTGCTGCGTTCTTCATCGATGC) (e.g., Smith and Peay, 2014; https://earthmicrobiome.org/protocols-and-standards/its/). Bacterial 16S rRNA genes were amplified with primers 799F and 1193R (CS1_799F: ACACTGACGACATGGTTCTACAAACMGGATTAGATACCCKG; CS2_1193R: TACGGTAGCAGAGACTTGGTCTACGTCATCCCCACCTTCC) (e.g., Vincent et al. 2022). First-stage PCR amplifications were performed in 10-µL reactions in a 96-well plate, using repliQa HiFi ToughMix (Quantabio, Beverly, MA, USA). PCR conditions for both reactions were 98 °C for 2 min, followed by 28 cycles of 98 °C for 10 s, 52 °C for 1 s, and 68 °C for 1 s. Subsequently, a second PCR amplification was performed in 10-µL reactions in a 96-well plate, again using repliQa HiFi ToughMix. Each well received a separate primer pair with a unique 10-base barcode, obtained from the Access Array Barcode Library for Illumina (Fluidigm, San Francisco, CA, USA; Item # 100-4876). A 1-µL aliquot of PCR product from the first-stage amplification was used as the template for the second stage, without cleanup. Cycling conditions were 98 °C for 2 min, followed by 8 cycles of 98 °C for 10 s, 60 °C for 1 s, and 68 °C for 1 s. Libraries were then pooled and sequenced with a 15% phiX spike-in on an Illumina MiSeq sequencer employing V3 chemistry (2 x 300 base paired-end reads). Library preparation and sequencing were performed at the Genomics and Microbiome Core Facility at Rush University (Chicago, IL, USA).

### 2.4. Amplicon sequence analysis and statistics

To analyze the amplicon sequence data, FASTQ files were imported and processed using the QIIME2 pipeline (version 2019.7) [39]. Sequence quality was visualized using the demux summarize plugin. For denoising of demultiplexed data, we used DADA2 [40], and resulting amplicon sequence variants (ASVs) were used for taxonomy classification with the Silva database (version 132, Quast et al., 2013) for 16S, and the UNITE database (version 8.3; Abarenkov et al., 2010; Nilsson et al., 2019) for ITS. 16S and ITS MiSeq forward and reverse reads were trimmed and merged, the low-quality sequences were filtered out, and the merged reads were uploaded to QIIME2 [44, 45]. Illumina-sequenced amplicon mistakes were rectified using DADA2 [40]. The resulting taxonomic tables (Supplementary Files: Taxonomy 16S and Taxonomy ITS) were subsequently uploaded to MicrobiomeAnalyst (microbiomeanalyst.ca; Chong et al., 2020; Dhariwal et al., 2017) for downstream analyses and visualization. Features with a minimum count of less than 20 and variance lower than 20%, based on interquartile range, were removed, and total-sum scaling was performed for normalization. Rarefaction curves, alpha diversity (Shannon diversity index), and beta diversity [Bray–Curtis distance matrix based on principal coordinates analysis (PCoA;Metsalu and Vilo, 2015)] were generated. Differentially abundant taxa at the ASV level were analyzed using DESeq2 [49], with cut-off padj < 0.05 (Supplementary Files: DESeq2). Abundance tables for the order and class levels were extracted to calculate bar plots. To assess the relationship between the chemistry of the banana container soil and the microbiota, canonical correlation analysis (CCA) was conducted using Vegan R package version 3.4.4 [48]. Pairwise Pearson’s correlation was calculated using cor function in R. The correlation heatmap plot was generated using the ClusViz tool (https://biit.cs.ut.ee/clustvis/). Conover’s test with Benjamini– Hochberg false discovery rate correction was used for pairwise comparisons, while a t-test with Benjamini–Hochberg correction for multiple comparisons was used to analyze banana plant size [50].

## 3. Results

### 3.1. Development of soil fatigue

Our analysis of crop yield data in plantations in the Jordan Valley in Israel, performed between 2008 and 2014, indicated a gradual deterioration in productivity beginning shortly after planting (Fig. 2), with a drop from the initial average crop yield of 82 ton ha^−1^ in the first year to 55 ton ha^−1^ in the sixth year-a decrease of 32% (for additional information on banana soil fatigue in this region, see Or et al., 2023). Moreover, in our experimental containers (see experimental design in Fig. 1), banana plants exhibited distinct growth reduction in response to varying increasing soil age (second-stage banana plants; Fig. 3). Notably, when plants were grown in younger soil, we observed the highest leaf (number 3) area and plant height, indicative of vigorous growth and leaf development. Conversely, in older soils, banana plants exhibited reduced leaf area and height, suggesting a potential decline in growth as the soil ages. The leaf area of the plant decreased from 14 cm² to 4 cm² at its lowest level in 36-month-old soil, and plant height was reduced from 90 cm to 28 cm in 30-month-old soil, representing a 70% reduction for both measurements compared to soils in which no plants had been grown previously (soil age 0). Note that both plant height and leaf area showed a rapid decline mainly during the first year, whereas plants in soils aged 12– 42 months maintained a similar small size. Significantly larger sizes were only observed in plants grown in 0- and 6-month-old soils (the youngest soils). The soil fatigue phenomenon predominantly manifested itself in our system within the first 12 months after planting (Fig. 3), faster than in the field (Fig. 2).

**Fig. 2.**
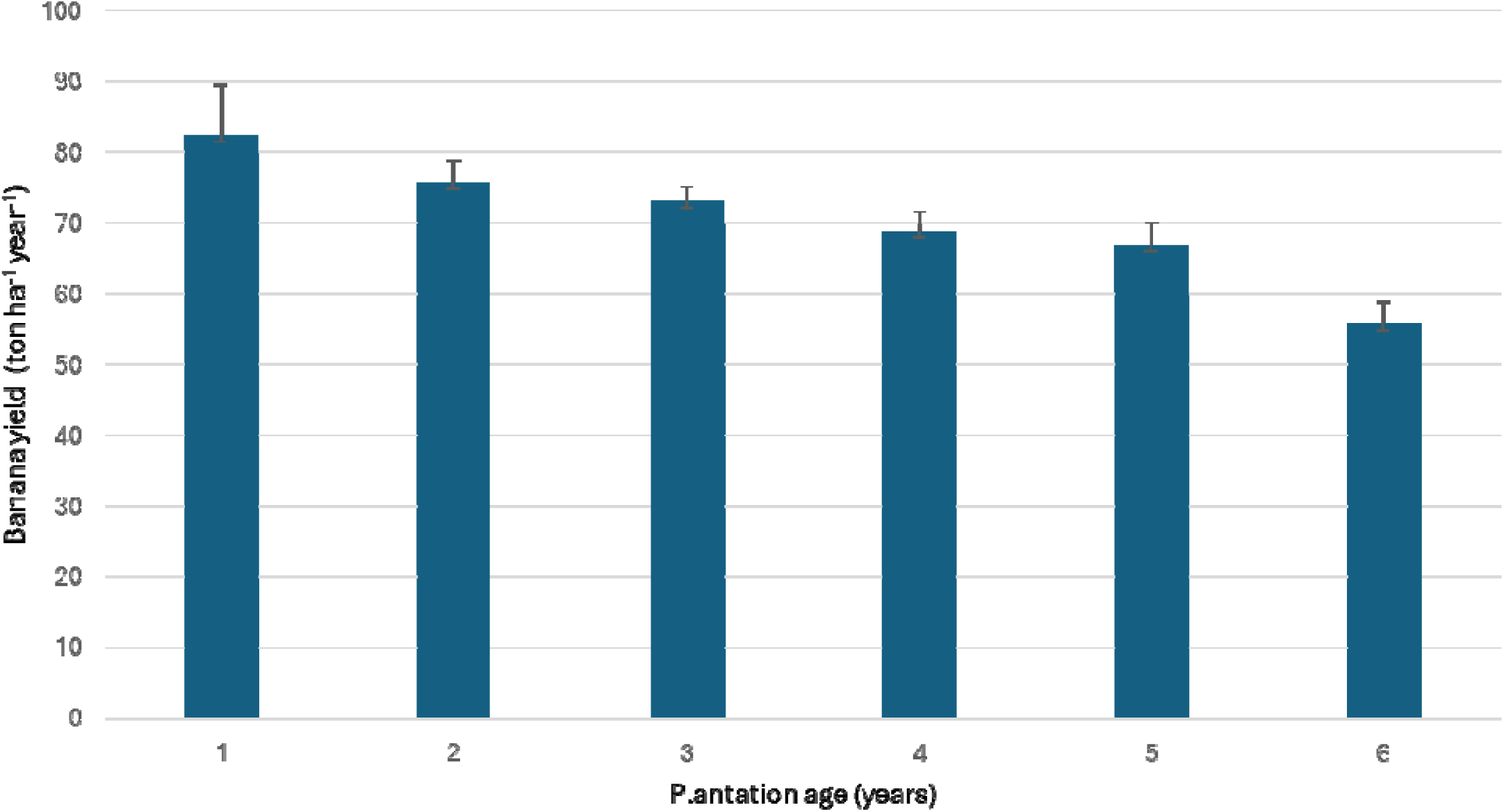
Decrease in annual banana yield per hectare, in commercial fields in northern Israel, over 6 years. Bars represent average yield in 10 commercial plantations, with standard error.

**Fig. 3.**
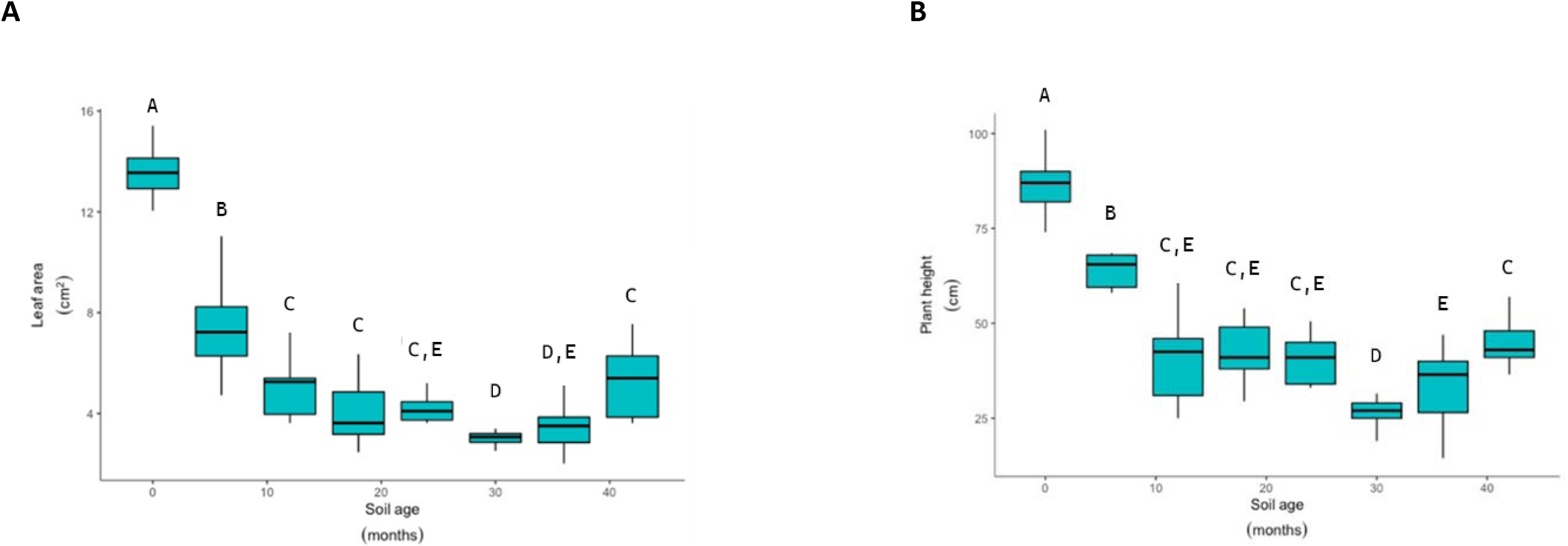
Effect of soil age on banana plant parameters. Boxplots depict the changes in (A) leaf number 3 area and (B) plant height in response to different primary(first stage) soil ages (0 to 42 months) in the containers. Different letters above the boxes indicate significant differences between groups (p < 0.05) based on t-test with Benjamini-Hochberg correction for multiple comparisons. Whiskers indicate range of observed values, n = 9.

### 3.2. Soil chemistry

To comprehensively explore the effect of soil age on its chemistry, soil samples were collected at the end of the 4 years of plant growth and analyzed. Soil samples were collected from each of three replicate containers of the specific soil age-0, 6, 18, 30, and 42 months-on the same date. A comprehensive suite of soil chemistry parameters was analyzed (Table 1 and Table S1). Data showed varied trends over time, with some analyses revealing remarkable stability, while others exhibited increasing or decreasing trends (Table 1). However, except for the decrease in pH level, none of the results for the following parameters were significant, due to high variability in values across the three replicates (Table S2). Potassium levels were three times higher in the youngest soil (0 months old) compared to the rest of the soils. There was a noticeable decrease in boron and phosphorus concentrations after 18 months. Concentration of available zinc, and available iron also increased throughout the period. TOC and UV, both markers of organic matter in the soil, behaved similarly, with an increasing over time The levels of the other elements-nitrate, ammonium, manganese, calcium & magnesium, chloride, sodium-and consequently, conductivity, were similar across soils of all ages. No abnormal salinization conditions developed, and the SAR range in the different soils represented a normal value for soils that have good structure and are generally favorable for plant growth.

**Table 1.**
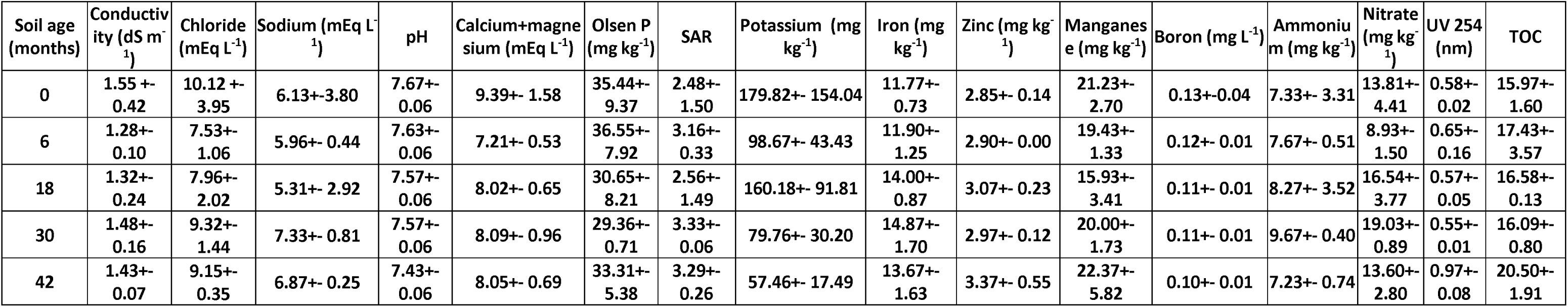
Effect of soil age on its chemical parameters. Mean values and standard deviations (n = 3) are presented for soil chemical parameters. Parameters were analyzed for soils aged 0, 6, 18, 30, and 42 months. TOC, total organic carbon; SAR, sodium adsorption ratio.

| Soil age (months) | Conductivity (dS m<sup>-1</sup>) | Chloride (mEq L<sup>-1</sup>) | Sodium (mEq L<sup>-1</sup>) | pH | Calcium+magnesium (mEq L<sup>-1</sup>) | Olsen P (mg kg<sup>-1</sup>) | SAR | Potassium (mg kg<sup>-1</sup>) | Iron (mg kg<sup>-1</sup>) | Zinc (mg kg<sup>-1</sup>) | Manganese (mg kg<sup>-1</sup>) | Boron (mg L<sup>-1</sup>) | Ammonium (mg kg<sup>-1</sup>) | Nitrate (mg kg<sup>-1</sup>) | UV 254 (nm) | TOC |
| --- | --- | --- | --- | --- | --- | --- | --- | --- | --- | --- | --- | --- | --- | --- | --- | --- |
| 0 | 1.55 +- 0.42 | 10.12 +- 3.95 | 6.13+- 3.80 | 7.67+- 0.06 | 9.39+- 1.58 | 35.44+- 9.37 | 2.48+- 1.50 | 179.82+- 154.04 | 11.77+- 0.73 | 2.85+- 0.14 | 21.23+- 2.70 | 0.13+- 0.04 | 7.33+- 3.31 | 13.81+- 4.41 | 0.58+- 0.02 | 15.97+- 1.60 |
| 6 | 1.28+- 0.10 | 7.53+- 1.06 | 5.96+- 0.44 | 7.63+- 0.06 | 7.21+- 0.53 | 36.55+- 7.92 | 3.16+- 0.33 | 98.67+- 43.43 | 11.90+- 1.25 | 2.90+- 0.00 | 19.43+- 1.33 | 0.12+- 0.01 | 7.67+- 0.51 | 8.93+- 1.50 | 0.65+- 0.16 | 17.43+- 3.57 |
| 18 | 1.32+- 0.24 | 7.96+- 2.02 | 5.31+- 2.92 | 7.57+- 0.06 | 8.02+- 0.65 | 30.65+- 8.21 | 2.56+- 1.49 | 160.18+- 91.81 | 14.00+- 0.87 | 3.07+- 0.23 | 15.93+- 3.41 | 0.11+- 0.01 | 8.27+- 3.52 | 16.54+- 3.77 | 0.57+- 0.05 | 16.58+- 0.13 |
| 30 | 1.48+- 0.16 | 9.32+- 1.44 | 7.33+- 0.81 | 7.57+- 0.06 | 8.09+- 0.96 | 29.36+- 0.71 | 3.33+- 0.06 | 79.76+- 30.20 | 14.87+- 1.70 | 2.97+- 0.12 | 20.00+- 1.73 | 0.11+- 0.01 | 9.67+- 0.40 | 19.03+- 0.89 | 0.55+- 0.01 | 16.09+- 0.80 |
| 42 | 1.43+- 0.07 | 9.15+- 0.35 | 6.87+- 0.25 | 7.43+- 0.06 | 8.05+- 0.69 | 33.31+- 5.38 | 3.29+- 0.26 | 57.46+- 17.49 | 13.67+- 1.63 | 3.37+- 0.55 | 22.37+- 5.82 | 0.10+- 0.01 | 7.23+- 0.74 | 13.60+- 2.80 | 0.97+- 0.08 | 20.50+- 1.91 |

### 3.3. Sequence analysis

Amplicon sequencing provided deep insights into the dynamics of the taxonomic composition of bacterial and fungal communities inhabiting the banana roots and agricultural soils of different ages during the development of soil fatigue. After processing the raw data, 3,284,253 bacterial and 1,196,826 fungal sequences were assigned to 11,565 and 1,080 ASVs, respectively. The numbers of sequences per sample ranged between 37,842 and 55,929 for bacteria, and between 5,912 and 24,521 for fungi, with an average of 46,257 and 16,857, respectively.

### 3.4. Niche effect on microbial communities

When communities from containers of different ages were analyzed together, there was no significant difference in alpha or beta diversities between rhizosphere and bulk soils (close and far from the root, respectively) for either 16S or ITS sequences (Fig. 4, Fig. S1). These findings implied that, in terms of Shannon alpha diversity and beta diversity, soil in these container systems could be considered a uniform entity, regardless of its spatial proximity to the roots. This, then, suggested that in this closed container and banana plant root systems, both soils are rhizospheric in nature. Therefore, roots and both soil samples were analyzed as two independent entities. As expected, significant differences were observed between communities from the roots and both soil types (Table S3): (i) alpha diversity in both the bacterial (Fig. 4A) and fungal (Fig. 4B) communities showed distinct differences between soil and root environments, with lower diversity in the latter community, and (ii) for beta diversity, a clear distinction in communities was observed between soils and roots for both bacteria (Fig. 4C) and fungi (Fig. 4D).

**Fig. 4.**
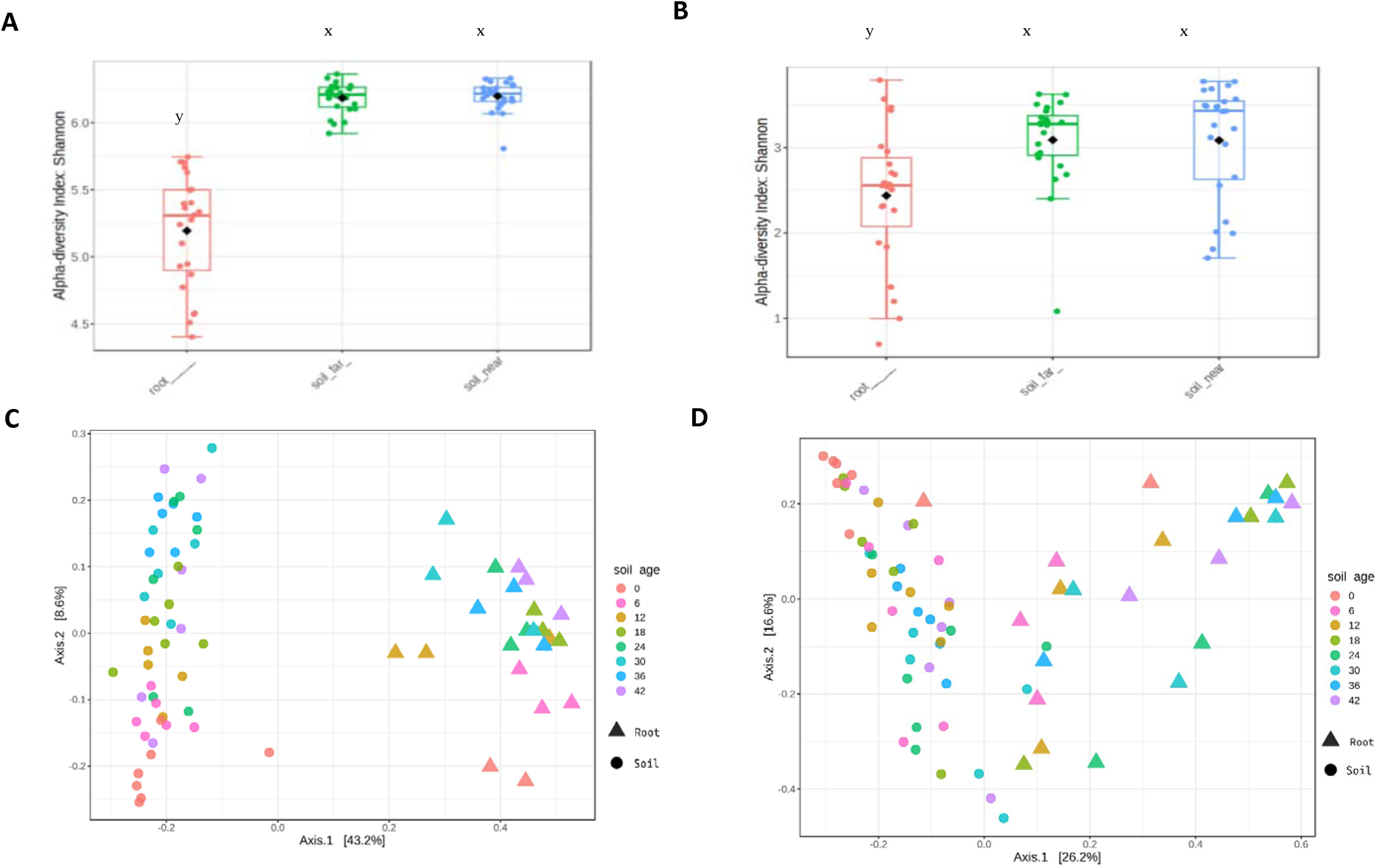
Microbial community analyses. Microbial diversity dynamics in roots and soil based on Bray–Curtis distances of (A) alpha diversity of bacterial communities (16S rRNA sequences), and (B) alpha diversity of fungal communities (ITS sequences). y, x indicate significant differences by t-test. Principal coordinates analysis plots of microbial community composition based on Bray–Curtis dissimilarity for 16S (C) and ITS (D) communities across different soil ages (0 to 42 months). Each point represents a microbial community sample, colors indicate soil age, and shapes indicate sample type (circle for soil, triangle for root).

### 3.5. Soil age effects on bacterial and fungal communities

PCoA based on the Bray–Curtis distance metric of 16S sequences was used to analyze sequence data, in order to evaluate the impact of soil age on beta diversity (Fig. S2). The ordination graph revealed clustering of eight groups associated with soil age, when visualizing the two-dimensional representation of the ASV matrix in both root (Fig. S2A) and soil (Fig. S2B) analyses. The results indicated a significant influence of soil age on most bacterial communities (*p* < 0.05) (Table S4). Notably, communities from soils aged 0 and 6 months were discernibly segregated from each other and from soils of other ages. As the soil analysis combined both samples (far from and close to the roots), the number of samples was double that for roots, enabling a determination of statistical significance. In the root, however, although there was a clear trend of separation of communities in root samples from 0 and 6 month soils, and from those in all other soil ages (Fig. S2A), it did not pass the stringent requirement for statistical significance.

The results of fungal ITS amplicon sequencing of soil samples were subjected to a similar analysis (Fig. S3). We found a much less significant influence of soil age on fungal communities (*p* < 0.05) (Table S5). Use of pairwise comparisons did reveal significant disparities in fungal communities between time 0 samples and all subsequent soil ages (Fig. S3B). In addition, significant discrepancies were apparent when comparing fungal communities in soil aged 36 months to those in soil aged 12 and 18 months. No significant differences were detected for the other soil ages. In the roots, as with the bacterial communities, a clear trend of separation of samples from soils aged 0 and 6 months, and from these compared to all other ages, was obvious (Fig. S3A), though not passing the stringent requirement for statistical significance.

To better understand the microbial community changes in the soil during the first year (the period when plant morphology was most damaged; Fig. 3), we categorized the samples into three distinct groups: soil at 0 months (youngest), soil at 6 months (young), and the remaining soil ages (old), and reanalyzed the subset data. This categorization aimed to provide a clearer visualization of the alterations occurring in that critical initial year for plants. There was a strong separation in the microbial PCoA plot between the three soil microbiomes of the different groups in both ITS (Fig. 5C) and 16S (Fig. 5D) analyses. All bacterial communities had significantly different Bray–Curtis index pairwise values (Table S4) (PERMANOVA, *p* < 0.05) for youngest vs. young (*p* = 0.002), youngest vs. old (*p* = 0.015), and young vs. old (*p* = 0.015) soils. Fungal communities were significantly different between youngest vs. young (*p* = 0.045) and youngest vs. old (*p* = 0.003) soils, but not young vs. old soils (*p* = 0.196) (Table S5). According to Kruskal–Wallis test, there was a significant difference in Shannon alpha diversity of the bacterial community between the youngest and young soils (*p* = 0.0032) and also between the youngest and old soils (*p* = 0.0032), but not for the young vs. old soils (*p* = 0.538) (Fig. 5). There was no significant difference (*p* > 0.05) in Shannon alpha diversity of the fungal community between the youngest, young, and old soils (Fig. 5). From this analysis, it could be inferred that the phenomenon of soil fatigue predominantly occurs in the first year, specifically between 0 and 6 months and between 6 and 12 months, aligning with our plant data (Fig. 3A, B).

**Fig. 5.**
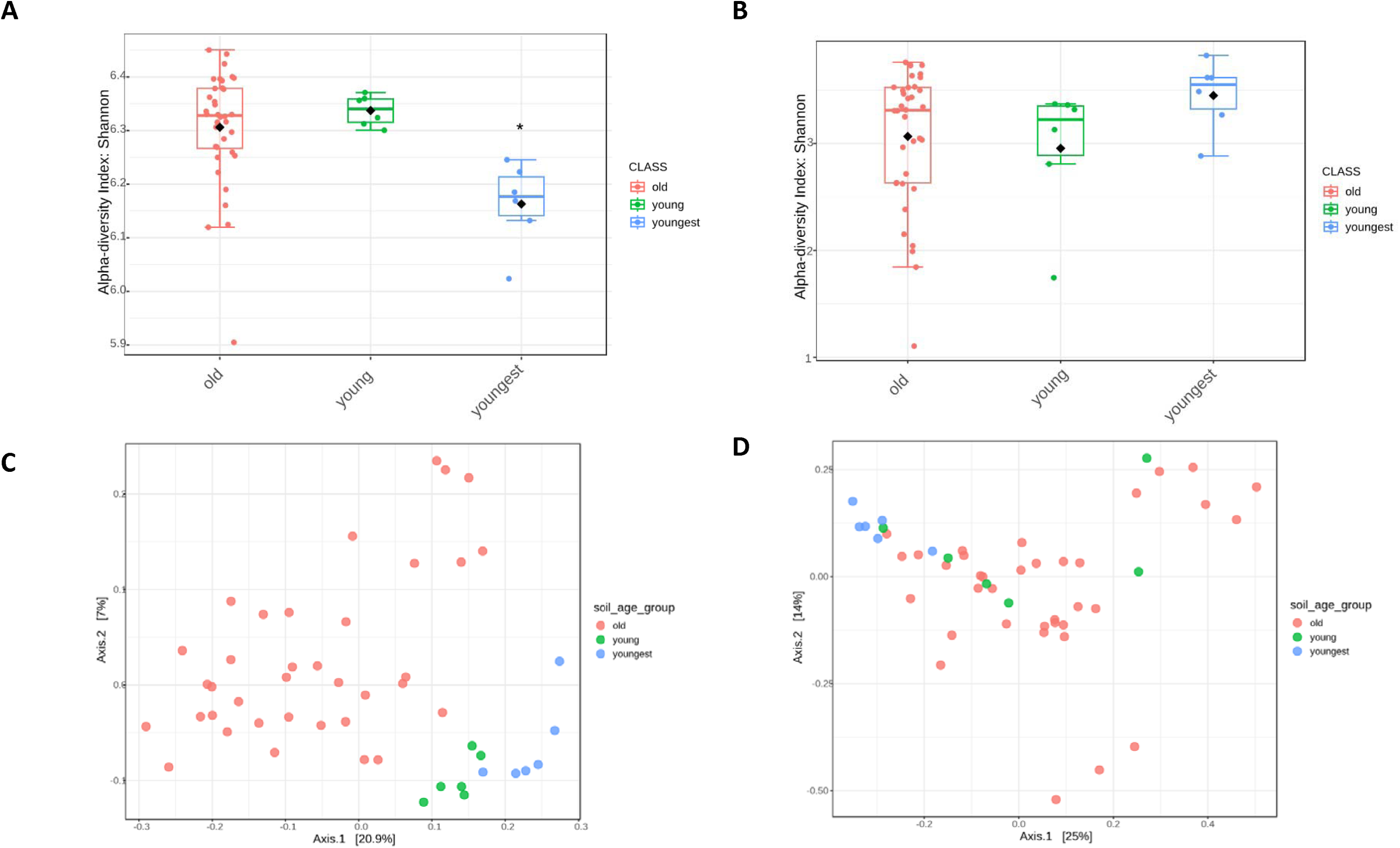
Microbial community analyses. Microbial diversity dynamics in soil samples. The group was separated into three subgroups based on Bray–Curtis distances of (A) alpha diversity of bacterial communities (16S rRNA sequences), and (B) alpha diversity of fungal communities (ITS sequences). Principal coordinates analysis plots of sequences of(C) 16S and (D) ITS communities. Diversity of microbial communities in rhizosphere with soil at 0 month (blue), 6 months (green), and all older soils (red). *Statistically significant difference (at p < 0,05).

Although the soil and root bacterial communities differed with soil age when ASV-based full communities were analyzed by PCoA (Fig. 5A, Fig. S2), when the 20 most dominant taxa were analyzed, even at the family and genus levels, only minor changes were noted (Fig. 6). When plants grew in age 0 soils, the families with the highest relative abundance in the roots were *Comamonadaceae* (29.33%) and *Sphingomonadaceae* (9.06%), and in the soil, *Nitrosomonadaceae* (14%) and *Gemmatimonadaceae* (5.8%). When plants were grown in soils of older ages, *Comamonadaceae* (relative abundance of 23.3% to 30.2%) and *Nitrosomonadaceae* (10.8% to 14.3%) remained the most dominant families in the root and soil communities, respectively.

**Fig. 6.**
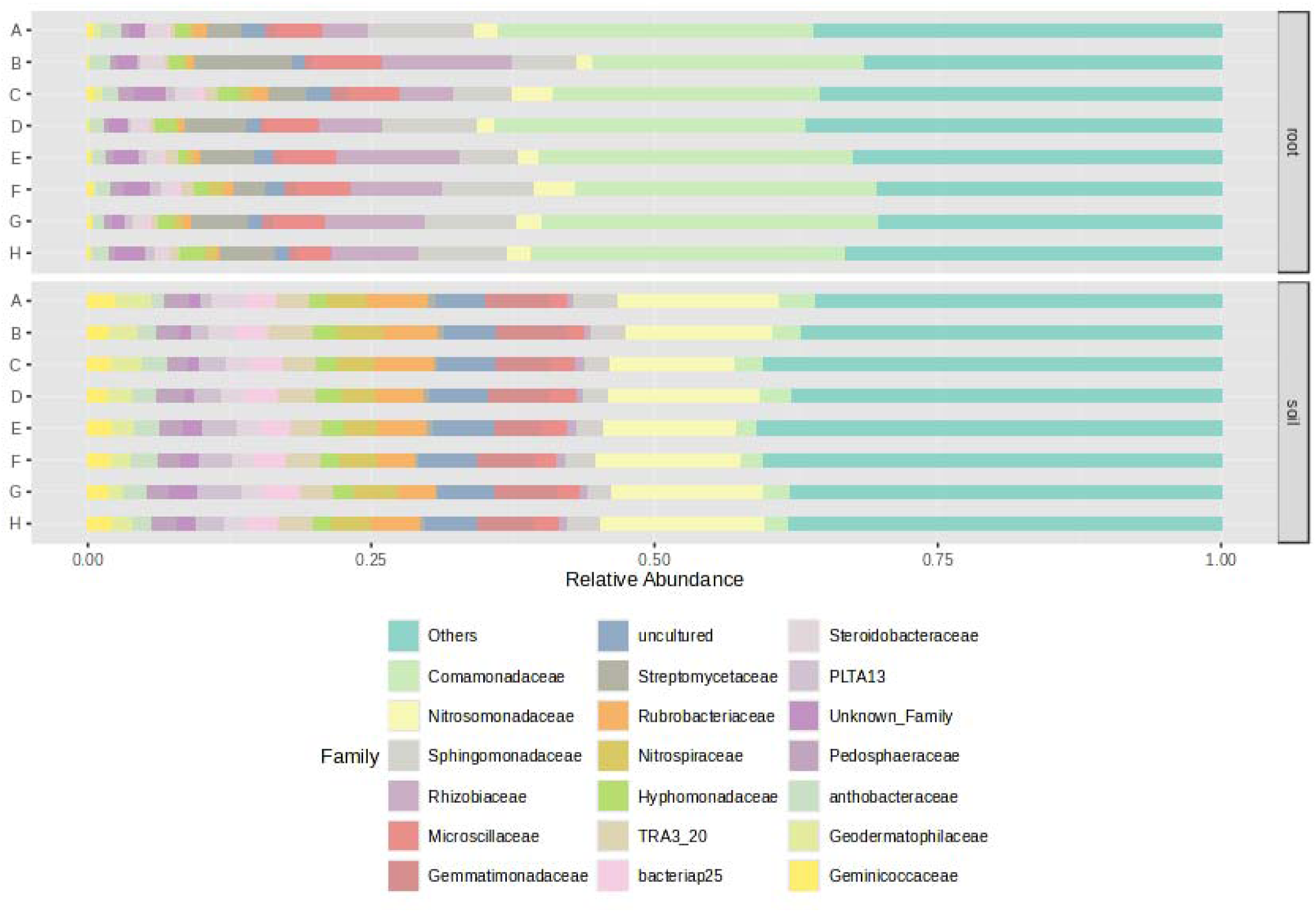
Family-level distribution of 20 top bacterial taxa in roots and soil. A-H: soil samples at different soil ages (A = 0 months, B = 6 months, C = 12 months, D = 18 months, E = 24 months, F = 30 months, G = 36 months, H = 42 months). Colored blocks represent individual families.

Interestingly, unlike the situation in bacterial communities, when analyzing the fungal ones via PCoA of the ITS sequences, only small changes could be detected (Fig. 4D, Fig. S3), whereas when looking at the relative abundance of the most dominant taxa, community changes were more visible (Fig. 7).

**Fig. 7.**
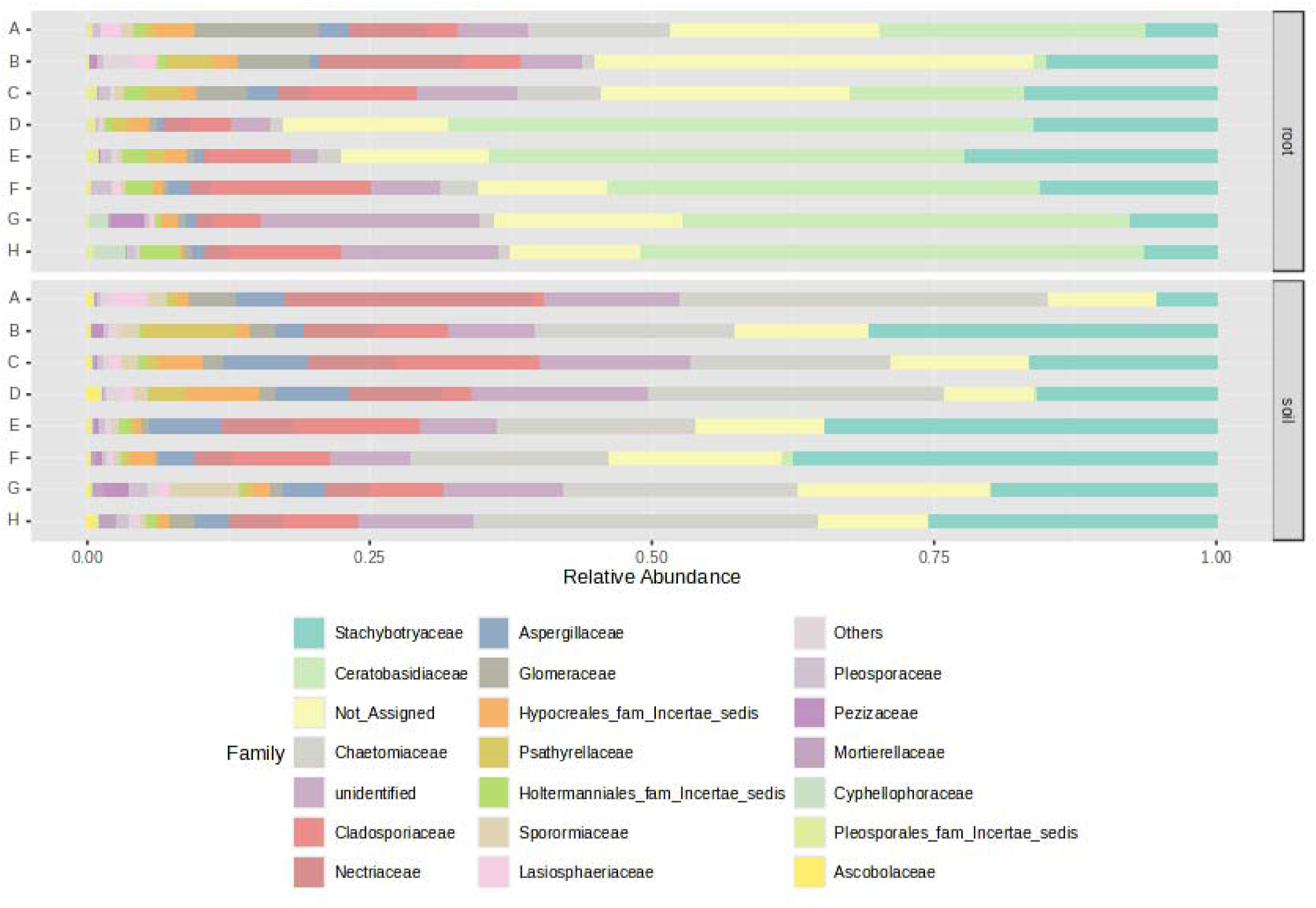
Family-level distribution of 20 top fungal taxa in roots and soil. A-H: soil samples at different soil ages (A = 0 months, B = 6 months, C = 12 months, D = 18 months, E = 24 months, F = 30 months, G = 36 months, H = 42 months). Colored blocks represent individual families.

In roots, the dominant fungal families were *Ceratobasidiaceae* (represented by the ASV related to *Rhizoctonia)*, with an average relative abundance of 1.3% increasing with soil age to 52%, followed by *Stachybotryaceae* (6.2% increasing with soil age to 22%). Albeit less dominant, *Cladosporiaceae* also exhibited a tendency to increase in relative abundance over time. *Nectriaceae, Chaetomiaceae, Psathyrellaceae, Lasiosphaeriaceae, Glomeraceae* and order Hypocreales exhibited a tendency to decrease with soil age.

In soil, the dominant fungal families were *Chaetomiaceae* (relative abundance increasing with soil age from 17.7% to 32.6%) and *Stachybotryaceae* (5.2% to 37.5%). Less dominant, but also exhibiting a tendency to increase over time were *Cladosporiaceae, Aspergillaceae, Pezizaceae, Mortierellaceae*, and *Pleosporaceae*. *Nectriaceae, Chaetomiaceae, Lasiosphaeriaceae*, and *Glomeraceae* exhibited a tendency to decrease in relative abundance with soil age. *Hypocreales* increased up to 18 months to a relative abundance of 6.53%, then decreased again to around 1.5% at 42 months.

### 3.6. Analysis of taxa with dynamic changes in relative abundance

A detailed analysis of root and soil microbiome dynamics across samples collected from containers representing different soil ages was performed and visualized as a heatmap (Fig. 8). Using the DESeq2 statistical framework, temporal shifts in microbial composition were identified. Some ASVs exhibited oscillatory patterns, with abundances fluctuating over the 42-month experiment, whereas others displayed unidirectional trends-consistently increasing or decreasing with soil age.

**Fig. 8.**
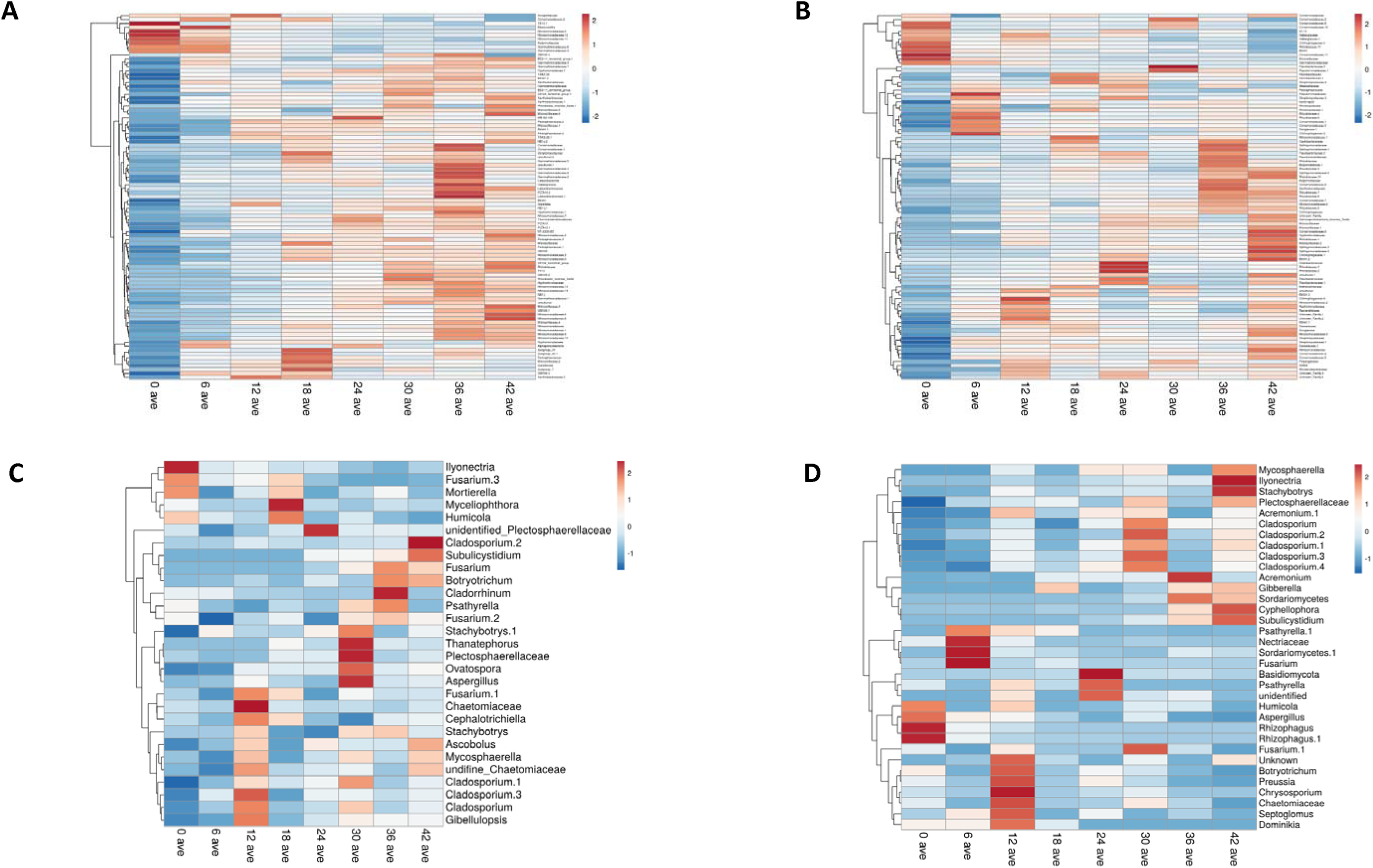
Differential abundance of bacterial and fungal taxa in the rhizosphere soil samples as determined by DESeq2. Heatmap and hierarchical cluster analysis using Euclidean distance and Ward linkage clustering algorithm at ASV level based on the relative abundances of biomarker taxa from (A) root (16S), (B) soil (16S), (C) root (ITS), and (D) soil (ITS). Color gradient from blue to red indicates relative abundance from low to high.

For the bacterial populations (Fig. 8 A, B), a significant increase in the relative abundance of certain bacterial family-level taxa was observed, particularly during the first year of the study. Notable taxa included families of *Flavobacteriaceae, Pedosphaeraceae, Rhizobiaceae, Latescibacteraceae, Pseudomonaceae, Haliangiaceae*, NB1-J (belonging to Nitrospirota), PLTA13 (belonging to Gammaproteobacteria), *Hyphomonadaceae, Paenibacillaceae*, and Blrii41 (belonging to Myxococcota) and order Acidibacter. Some taxa that were not well represented (less than 1% relative abundance), such as A0839 (belonging to Rhizobiales), *Thermoanaerobaculia, Hyphomicrobiaceae, Pleomorphomonadaceae*, and subgroup 2, 22, 10 (belonging to Acidobacteriota) were also identified (Fig. 8A, B and Supplementary Files: RA increasing 16S). Conversely, a decrease in some bacterial taxa was noted over time, again particularly during the first year. These included *Xanthobacteraceae, Moraxellaceae, Devosiaceae, Gemmatimonadaceae, Sphingomonadaceae*, and *TK* 10 (belonging to Chloroflexi). Here, again, some taxa that were not well represented (less than 1% relative abundance), such as S0134 and Bd2-11 (both belonging to Gemmatimonadota), *Beijerinckiaceae, Acidobacteriota* subgroup 7, and *Azospirillaceae*, were identified (Supplementary Files: RA decreasing 16S). These observations indicated that the more pronounced shifts occur within the initial months of the experiment, which is in correlation with the results of the beta diversity PCoA results (Fig. 4C, Fig. S2). A similar trend in abundance shifts was noted in the fungal populations (Fig. 4D, Fig. S3).

Interestingly, an increase in the relative abundance of several fungal taxa that are often associated with pathogenic groups was observed (Supplementary Files: RA increasing ITS). These included members of the *Cladosporiaceae, Trichocomaceae, Stachybotryaceae, Aspergillaceae, Plectosphaerellaceae*, and *Ceratobasidiaceae* (mainly a *Rhizoctonia* ASV). In contrast, taxa associated with symbiotic or beneficial relationships, including *Glomeraceae* (*Rhizophagus* among them), *Lasiosphaeriaceae, Chaetomiaceae*, and members of the *Nectriaceae*, exhibited a declining trend (Fig. 8C, D and Supplementary Files: RA decreasing ITS).

These findings underscored the significant microbial community shifts occurring during the soil fatigue phenomenon, which were consistent with the beta diversity analyses.

### 3.7. Soil chemistry is correlated with shifts in microbial communities

To better understand the relationships between microorganisms and the changing environmental conditions in the banana rhizosphere, soil chemistry parameters were analyzed together with community parameters using CCA (Supplementary Files: CCA chemistry correlation 16S, CCA chemistry correlation ITS). The CCA graph visually represents the relationships between environmental variables and ASVs with respect to soil age (Fig. 9). We selected 14 parameters [pH, conductivity, chloride, sodium, calcium & magnesium, potassium, iron, zinc, manganese, boron, ammonium, organic matter (TOC), phosphorus (Olsen P), and SAR] for CCA based on significance tests (see section 2). Several key environmental variables were found to be correlated with bacterial and fungal communities. Potassium and pH exhibited a strong positive association with microbial community composition, whereas TOC and iron vectors were oriented with a negative correlation. As indicated by the arrow lengths of the environmental variables in the CCA biplot, the strongest correlations with the bacterial and fungal community composition were for pH, potassium, TOC, and iron. For example, high correlations were observed between the parameters pH and potassium and bacterial ASVs related to *Comamonadaceae* and OM190 (phylum Planctomycetota), while a strong correlation was found between SAR and TOC and several ASVs related to *Latescibacteraceae*. High correlations were also observed between boron and phosphorus (Olsen P) parameters and bacterial ASVs related to *Blastocatellia, Azospirillaceae, Beijerinckiaceae*, and *TK* 10 (phylum Chloroflexi), as well as several ASVs of *Gemmatimonadaceae* and *Nitrosomonadaceae* (Fig. 9A). Regarding fungal communities (Fig. 9B), there was a correlation between pH and ASVs related to *Chaetomiaceae* and *Acremonium*, between potassium and ASVs related to *Humicola* and *Fusarium*, and a strong correlation between TOC and zinc and the ASVs related to *Alternaria, Subulicystidium, Chaetomium, Mortierella*, and two ASVs related to *Mycosphaerella*. In all cases, most 16S and ITS ASVs clustered in association with soil ages 30 and 42 months. To better understand changes in the bacterial and fungal populations in correlation with soil chemical parameters, a heatmap analysis was employed based on sequences whose relative abundance shifted significantly in correlation with the soil parameters (Fig. 10). The results revealed the existence of three major distinct groups among the soil parameters to which the microorganisms were correlated (labeled I, II, and III). These organisms correlated to the soil parameters in two models (X and Y for bacteria and x and y for fungi). In the bacteria, group I included calcium & magnesium, chloride, conductivity, and manganese, group II contained nitrate, TOC, UV, zinc, iron, SAR, and sodium, and group III contained boron, pH, phosphorus (Olsen P), potassium, and ammonium. In the fungi, the grouping was similar, with group I including nitrate, ammonium, conductivity, and chloride, group II containing TOC, UV, zinc, iron, SAR, and sodium, and group III comprising calcium & magnesium, boron, pH, phosphorus (Olsen P), and potassium. These findings highlight distinct clusters of soil parameters driving microbial community composition in the system.

**Fig. 9.**
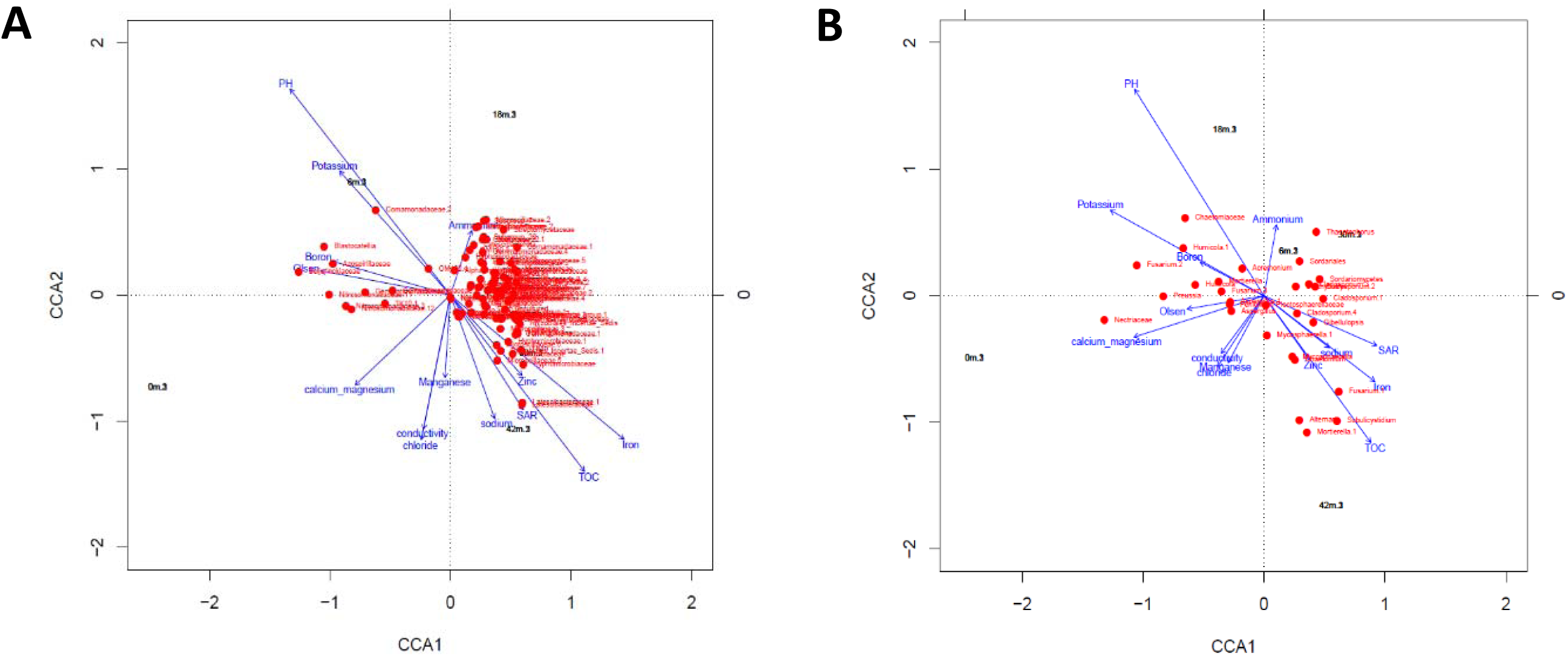
Canonical correspondence analysis (CCA) biplot of species and environmental variables. This biplot illustrates the relationships between environmental variables (blue arrows) and species (red dots), based on the CCA for sequences of 16S (A) and ITS (B). The black text represents soil age, showing its distribution along the environmental gradients. Arrows indicate the direction and magnitude of soil chemical parameters correlated with microbial community structures.

**Fig. 10.**
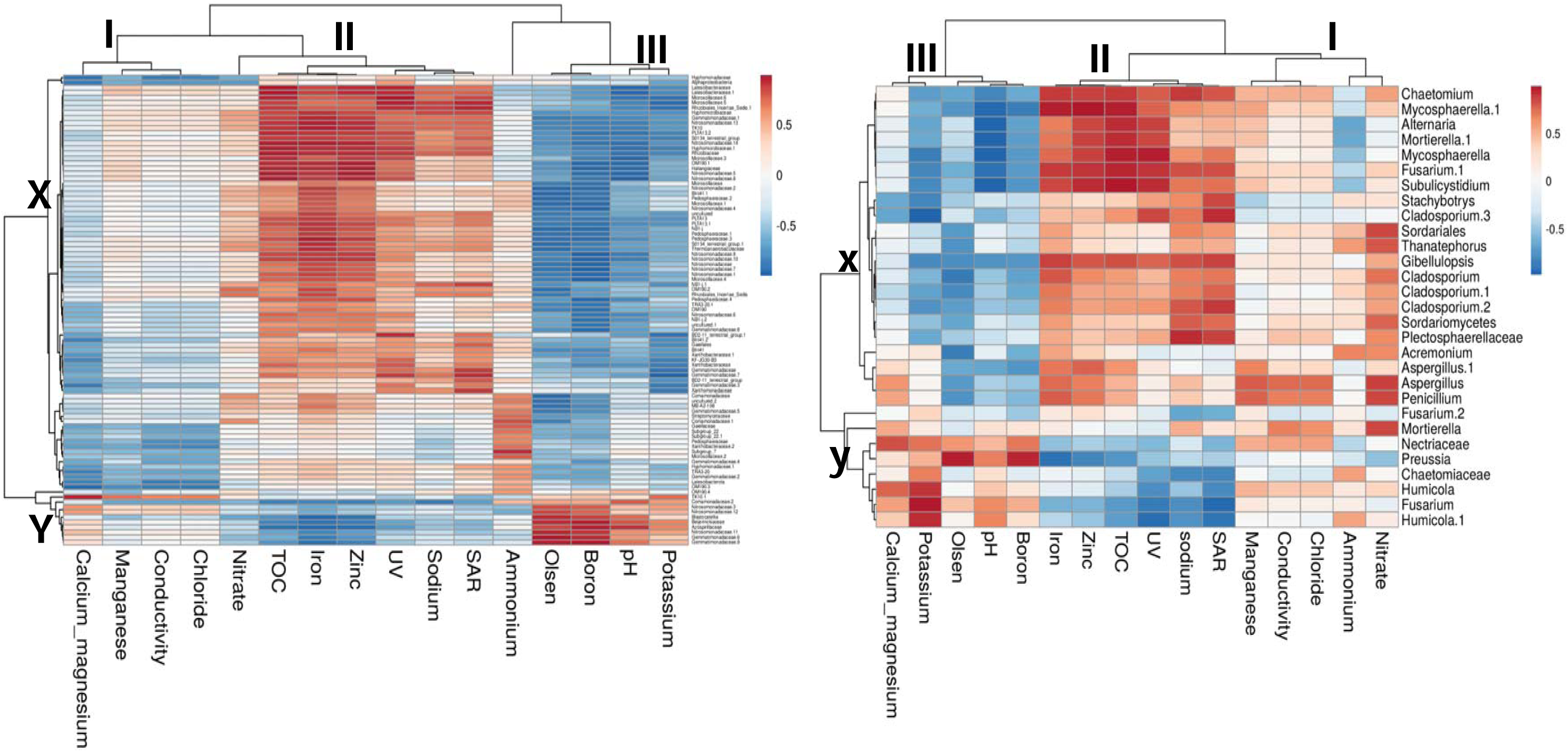
Heatmap analysis of microbial ASVs’ correlation with soil chemistry. Heatmap and hierarchical cluster analysis using Euclidean distance and Ward linkage clustering algorithm at ASV level, based on the relative abundances of biomarker taxa for (A) 16S and (B) ITS clustering, are applied on both axes, with color intensity representing the correlation of each taxon (X, Y for 16S and x, y for ITS) with different soil chemical parameters (I, II and III). Red indicates positive correlation; blue denotes negative correlation.

## 4. Discussion

The phenomenon of soil fatigue is characterized by a reduction in plant growth with cultivation time. Its detrimental impact on banana yield challenges the sector’s profitability. When yield falls below the economic viability threshold (about 50 ton ha^−1^ year^−1^), generally about 8 to 12 years after planting, growers are forced to uproot their entire field. Despite the economic burden posed by soil fatigue, our understanding of its underlying causes remains limited.

In a large survey conducted in 10 commercial plantations in the Jordan Valley over a period of 6 years, the effect of soil fatigue was reflected in a 32% drop in profitability (Fig. 2). To investigate the root and soil microbial and chemical parameters involved in this phenomenon, we developed a controlled banana growth system that incorporated soils of varying ages (Figs. 1 and 3). Banana plants of identical age and size were cultivated in soils previously used in banana cultivation for different durations. Remarkably, plant growth varied significantly with soil age (Fig. 3). The larger plants were observed in soils in which bananas had not been previously cultivated, whereas the smallest and least developed plants were found in soils with a history of banana cultivation for 18 months or longer (soil ages of 18–36 months). These findings align with previous studies indicating the importance of soil age in plant growth [31, 51, 52].

Rhizosphere microorganisms play a crucial role in facilitating the growth and development of crops by engaging in complex interactions and mutualistic relationships within the root system. In addition, plants, being sessile organisms, constantly face environmental stresses; they can mitigate these pressures by harnessing indigenous microbiomes for their defense mechanisms [53, 54]. Our research focused on banana soil and root microbiology, utilizing amplicon sequencing of the bacterial and fungal communities (Figs. 4–7), as well as soil chemistry (Table 1, Figs. 9, 10). This approach aligns with previous research on soil fatigue, which has been explored from physical [55], chemical [30], and biological perspectives, particularly regarding the emergence of pathogens and the associated decline in biodiversity [56, 57]. As expected from previous studies on other plant and soil microbiomes, the root communities in our system were always different from those of the soil and rhizosphere, typically with lower alpha diversity and clustering separately from those of the soil and rhizosphere [58, 59] (Fig. 4). In our experiment, the alpha and beta diversities in bulk soil and the rhizosphere were unexpectedly similar; based on previous studies, we would have anticipated greater diversity in the former relative to the latter [60]. This observation is likely because our experiment was conducted in containers rather than directly in the soil, which may have allowed the roots’ influence to extend into the bulk soil. Thus, we hypothesize that in our experimental system, the “bulk” soil is actually also rhizospheric soil.

Bacterial communities in both the soil and roots exhibited more pronounced overall temporal changes compared to the fungal communities (Figs. S2 and S3, respectively). We observed distinct bacterial clusters, with the clearest separations occurring at soil ages of 0 and 6 months. Partial distinctions were also observed at 12 and 18 months, whereas clustering became less defined at older soil ages. This was true and significant for soil communities, as well as root communities (where the trend was clear but due to low replicate numbers, lacked statistical significance). For the ITS analysis, the trend was similar, although significance was only high in soils aged 0 and 6 months compared to the “older” soils.

Although the plants were all planted simultaneously (newly planted in all soil age treatments) and were identical to start with, their development was closely linked to soil age and microbial composition. This suggests that soil characteristics, shaped by the duration of prior plant growth, significantly influence microbial populations, including those associated with the roots of the planted bananas. Despite only minor but significant changes in pH (Table 1), the large shifts in microbial communities within the soil likely played a key role in driving the observed differences in plant growth and development.

Pairwise Bray–Curtis dissimilarity indices for the 16S sequences divided into youngest, young, and old samples (soil age 0, 6 months, and all later ages, respectively) were significantly different from each other (i.e., youngest vs. young, youngest vs. old, and young vs. old) (Fig. 5C). For ITS communities, significant differences were observed between the youngest and young, and youngest and old ages, but not between the young and old ages (Fig. 5D). In addition, according to the Kruskal–Wallis test, for the Shannon diversity index, the 16S communities showed significant differences between the youngest and young, and youngest and old soils, but not between the young and old soils (Fig. 5A), whereas there was no significant difference for ITS communities (Fig. 5B). All of these results align with previous studies that indicated that fungal communities, compared to bacterial ones, exhibit greater resilience to environmental changes. For example, bacteria were shown to hold a much bigger place than fungi in the dynamic changes taking place during forest restoration [61]. Moreover, bacterial communities were shown to be more susceptible than fungal ones to the process of forest succession [9, 62, 63]. The findings are also consistent with studies conducted by Brown and Jumpponen (2015) and Zhang et al. (2018), which indicated that fungal communities exhibit minimal changes over time and are mostly influenced by plant species. Other studies conducted on a variety of crops also found no significant variations in fungal diversity indices throughout different periods of continuous cropping [66–69]. It is well accepted that the faster reproduction rates and better metabolic flexibility of bacteria enable them to respond and adapt more rapidly than fungi to changes in environmental conditions [70–72]. Furthermore, the absence of any alterations in fungal community structure following the first time point and throughout the last 3 years of growth indicates that in our system, the fungal community also shifts at a slower rate than the bacterial one.

To better understand the composition of microbial communities involved in the soil fatigue phenomenon, we used microbiome analysis (https://www.microbiomeanalyst.ca) to focus on the changes in relative abundance of specific populations in relation to soil age. We found that the phyla Proteobacteria, Actinobacteria, Myxococcota, Gemmatimonadota, and Acidobacteriota dominated in the banana rhizosphere, and Proteobacteria, Actinobacteria, Myxococcota, Bacteroidota, and Firmicutes dominated the root bacteria. This fits with the results of other studies showing that Proteobacteria, Actinobacteria, and Acidobacteriota are major microbial community members in most soils [73, 74], and of studies showing that members of the Proteobacteria, Actinobacteria, Acidobacteria, Chloroflexi, Firmicutes, Bacteroidetes, and Gemmatimonadetes are dominant bacterial phyla in banana soil [27, 34]. The results also agree with studies showing that Proteobacteria, Acidobacteriota, Actinobacteriota, Bacteroidota, Firmicutes, and Gemmatimonadota are dominant in the banana rhizosphere [27, 34]. *Comamonadaceae* and *Nitrosomonadaceae* were found to be the most dominant families in soil and root communities, respectively (Fig. 6), similar to many agricultural systems [75–77]. Dominance of members of the family *Nitrosomonadaceae*, commonly involved in nitrification, could be explained by the fact that our system was consistently fertigated with ammonium-containing fertilizer. Ammonia oxidation by members of the *Nitrosomonadaceae* may also have contributed to the observed decrease in soil pH (Table 1).

DESeq2 statistical analysis was utilized to identify temporal fluctuations in the microbial composition at the ASV level. This revealed a notable rise in specific bacterial families (Fig. 8A, B). The noteworthy families included an unknown *Acidibacter* (belonging to Gammaproteobacteria), *Flavobacteriaceae, Hyphomonadaceae*, NB1-J (belonging to Deltaproteobacteria), *Paenibacillaceae, Pseudomonaceae, Haliangiaceae, Pedosphaeraceae*, PLTA13 (belonging to Gammaproteobacteria), and Blrii41 (belonging to Myxococcota). *Acidibacter* has been previously shown to play a key role in the breakdown of soil organic matter (Chen et al., 2020; Luo et al., 2023). Similarly, *Flavobacteriaceae, Latescibacteraceae*, and *Pedosphaeraceae* have been shown to be associated with organic matter decomposition and soil nutrient cycling [80–82]. The increases in the relative abundance of *Paenibacillaceae* and Blrii41 were continuous and more pronounced after the first year, indicating that these bacteria could be reacting to long-term variations in soil conditions, such as the increase in organic carbon levels, as described previously [83, 84]. The detected increase in soil organic matter following plant growth may explain the increase in relative abundance of these families.

Some studies [5, 85–88] have indicated a symbiotic relationship wherein plants under pathogenic stress recruit assistance from bacterial partners such as *Lasiosphaeriaceae, Flavobacterium, Chitinophagaceae*, and *Sphingomonas*, corresponding with the observed proliferation of these taxa in our system. This finding suggests a possible role for these bacteria in plant defense against pathogens (i.e., fungi), potentially involving the production of antimicrobial compounds or modulation of plant immune responses. The further observation of an increase in the relative abundance of predatory family Blrii41 [89, 90] in both our soil and root systems suggests a role in promoting dynamic changes in the community structure, nutrient cycling dynamics, and overall ecosystem stability.

In contrast, a decline in the relative abundance of certain families was observed over time, especially within the first year. These included members of the *Gemmatimonadaceae, Moraxellaceae, Sphingomonadaceae, Xanthobacteraceae, Azospirillaceae*, and the genus *Acidovorax* of the family *Comamonadaceae*. The family *Gemmatimonadaceae*, commonly found in habitats with low nutrient levels, may have faced competition when nutrient levels fluctuated [91, 92]. Some members of the *Acidovorax, Azospirillaceae, Moraxellaceae*, and *Xanthobacteraceae* are known to function as symbiotic bacteria, exerting beneficial effects on plants by producing secondary metabolites and hormones that promote plant growth (Brescia et al., 2023; Giordano et al., 2012; Han et al., 2016; Siani et al., 2021; Armanhi et al., 2018; Beirinckx et al., 2020; Benitez et al., 2017). The observed decline in potentially beneficial microorganisms may affect plant–pathogen interactions, promoting a decline in overall plant health and development, as seen in soil fatigue.

In the current study, Ascomycota was the most abundant fungal phylum in banana roots and soil over time, with a predominance of its classes *Sordamriomycetes* and *Dothideomycetes*, and *Agaricomycetes* of the Basidiomycota was highly dominant in the roots (Fig. S4). This correlates with a previous, thorough study of the core fungal microbiome in banana conducted by Birt et al. (2023).

An increase in the relative abundance of fungi belonging to the *Dothideomycetes* was observed with soil age in both soil and roots. These comprise a diverse group of fungi with varied ecological roles, including plant pathogens, endophytes, and saprobes [100–103]. A significant increase was observed in the relative abundance of several fungi belonging to families such as *Plectosphaerellaceae, Stachybotryaceae* and genus *Cladosporium* known to include pathogenic representatives [104–106], in both soils and roots during the initial year of the experiment (Fig. 7). Over a longer period, a relative increase in the abundance of fungi from the *Ceratobasidiaceae* (represented mainly by *Rhizoctonia* ASVs) was observed in the roots, and of *Trichocomaceae* and *Aspergillaceae* in both roots and soils. Both of these families are known to have pathogenic representatives [107–109]. Interestingly, the genera *Cladosporium, Stachybotrys* along with the family *Ceratobasidiaceae* have representatives that have been shown to be specific pathogens of banana plants (Chen et al., 2020; Samarakoon et al., 2021, 2021; Surridge et al., 2003).

In contrast, a noticeable decline in the occurrence of certain fungal families, such as *Glomeraceae* (including specifically the genus *Rhizophagus)*, *Chaetomiaceae*, and *Lasiosphaeriaceae*, which are typically found in plant growth-supporting symbiotic relationships with plant roots and have potential biocontrol activity [111–115] was noted. This trend suggests a dynamic shift in the fungal population structure, characterized by an increased relative abundance of suspected pathogenic species and decrease in that of possibly beneficial ones. These changes in microbial communities may theoretically be involved in the damage inflicted on the plants during the development of the soil fatigue phenomenon.

Soil pH was the most dominant factor influencing bacterial and fungal community compositions (Fig. 9) and was the only chemical element that changed significantly (Table S2), even though there was only a 0.3 pH unit variation between different soil ages (Tables S1 and S2). The results align with those of other studies showing that a slight variation in soil pH, even within a range of less than 0.5 units, can have a significant impact on the optimal growth of microbial communities in various cropping systems [117–120]. In fact, continuous ammonium-containing fertilization of fields (especially those containing ammonia, such as in our case) may causes an increase in the process of nitrification, resulting in the release of protons and a higher level of soil acidity [121–126]. In addition, some bacteria identified as key members in our system belong to groups that have also been shown to be associated with increased soil acidification, including *Acidibacter*, *Flavobacteriaceae*, and *Paenibacillaceae* (Xu et al., 2023; Zhang et al., 2022). This, may promote the observed changes in the microbial communities, including the significant proliferation and spread of soilborne pathogens, possibly associated with the development of soil fatigue [129, 130].

Organic matter measured by TOC and UV absorbance levels increased with soil age, indicating potential accumulation of organic matter and changes in soil organic carbon resulting from plant growth [131, 132]. These findings suggest that the continuous plant growth in our system leads to an increase in organic matter, consistent with previous studies in other crops showing a significant rise in soil nutrient levels and improvements in soil physicochemical characteristics as a result of prolonged and uninterrupted cultivation, leading to an increase in the population of microorganisms involved in soil organic matter production [119, 133, 134]. Our findings revealed that soil organic matter (TOC) is the second-most dominant factor affecting microbial communities (Fig. 9). A clear increase was noticed with time in the relative abundance of ASVs belonging to groups that have been previously shown to be associated with the accumulation of soil organic matter. These include families such as *Flavobacteriaceae, Rhodocyclaceae, Hyphomonadaceae*, and *Microscillaceae* [86, 135–137]. The increase in organic matter may support the growth of fungal groups known to be saprophytic and/or pathogenic (*Ceratobasidiaceae* in roots, and *Trichocomaceae* and *Stachybotryaceae* in roots and soils), which was noted with increasing soil age, and discussed above.

With soil age, a decline in the relative abundance of members of the *Gemmatimonadaceae* was correlated with diminishing available phosphorus (Olsen P) in the soil (Fig. 10). Some members of the genus *Gemmatimonas* are known for their ability to solubilize phosphorus [91, 92], thus the decline in their populations could contribute to the decrease in phosphorus availability in the soil.

Iron is a vital micronutrient required for key microbial metabolic processes, including respiration, DNA synthesis, and the production of virulence factors; thus, pathogenic fungi have developed diverse strategies to acquire and utilize it [138–140]. In our system, we observed a temporal increase in available iron levels in the soil, which may have affected the relative abundance and activity of fungal groups containing pathogenic members, potentially enhancing their pathogenic potential. These changes could be one of the soil parameters contributing to the onset of soil fatigue.

## 5. Conclusion

Little is known about the development of the banana plant root microbiome, especially in relation to soil fatigue. This study followed the evolution of chemical conditions, as well as bacterial and fungal populations, in soil and banana roots as soil fatigue developed. We observed a significant decrease in soil pH along with some trends in other parameters, including an increase in soil organic matter (TOC, UV) and available iron, coupled with decreases in phosphorus and potassium. These changes indicated complex interactions between soil chemistry and organic matter dynamics correlated with soil age, and associated with dynamic shifts in both bacterial and fungal communities. Significant shifts in microbial community composition included an increase in the relative abundance of bacterial families which are often associated with organic matter degradation. Conversely, bacterial families that are known to include plant-beneficial members exhibited a decline in relative abundance. These trends suggest a possible loss of bacterial diversity which would favor beneficial interactions and a proliferation of bacterial groups potentially linked to soil-degradation processes. In addition, although we did not observe an increase in the relative abundance of specific known banana pathogens, there was a rise in the relative abundance of several families that are known to contain pathogenic fungi, in parallel to a decrease in beneficial fungi, which could potentially lead to a decline in soil health and fertility, manifested as soil fatigue. The implications of this study are significant for the agricultural industry, justifying further research into the inoculation of beneficial fungi, such as *Rhizophagus* (the relative abundance of which decreased with soil age), plant growth-promoting bacteria, and biocontrol agents against potential soil pathogens, alongside improved fertilizer and pesticide management. Such research will increase knowledge that is most relevant to the development of effective banana microbiome-management practices and to better agricultural practice. Despite the extensive analyses presented in this article, including hypotheses on potential interactions, correlations between plant parameters, soil chemistry, and microbial community dynamics, and the application of various statistical methods, similar to the situation in other crops, it still remains impossible to definitively identify the specific factors that are directly responsible for soil fatigue in banana cultivation.

## Data availability statement

All data from this study are fully accessible in the public repository NCBI under bio-projects PRJNA1169519.

