## Supplementary material for "Dynamics in microbial communities associated with the development of soil fatigue in banana": Fig.S and Table S

### Slide 1
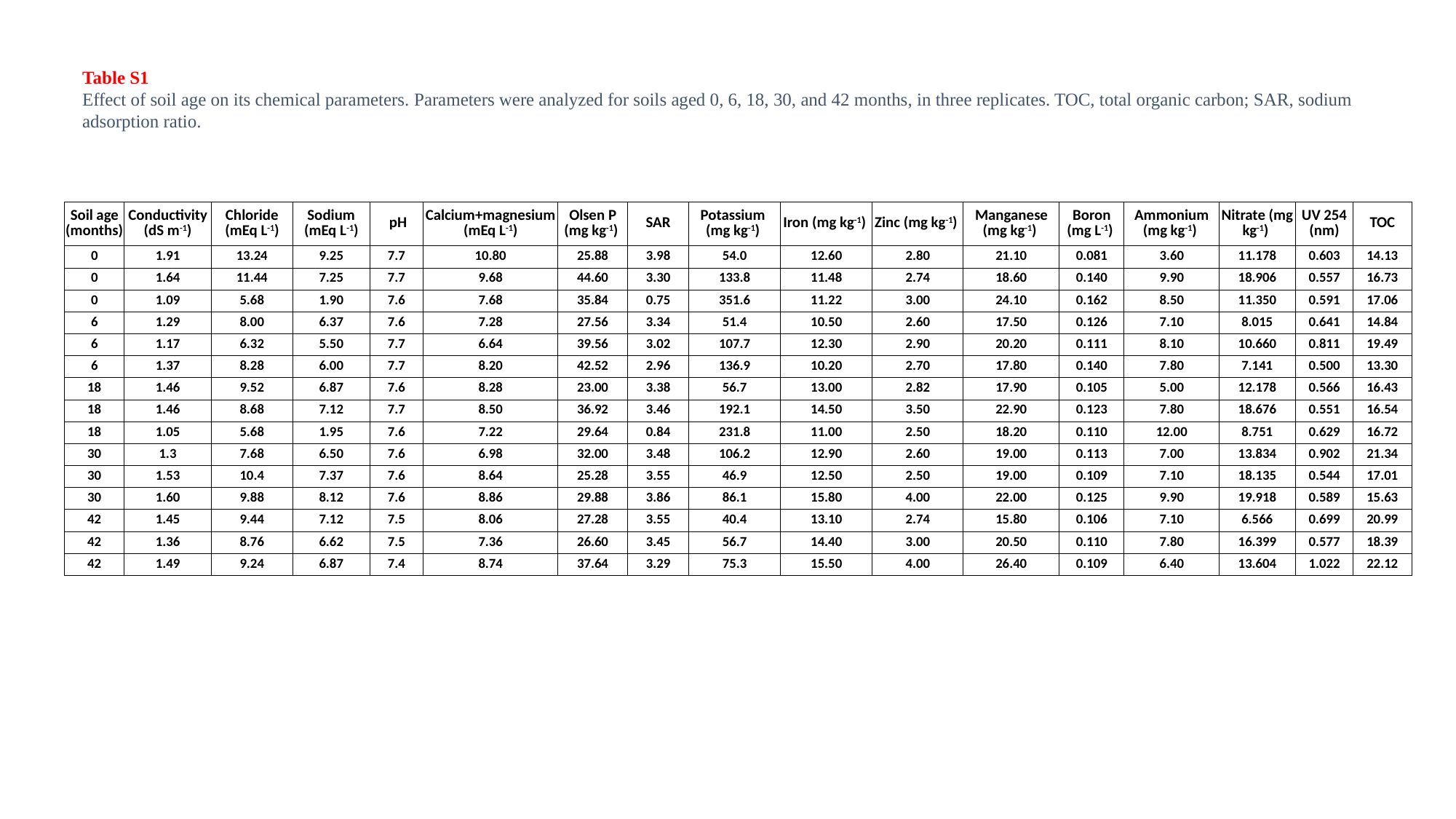

Table S1
Effect of soil age on its chemical parameters. Parameters were analyzed for soils aged 0, 6, 18, 30, and 42 months, in three replicates. TOC, total organic carbon; SAR, sodium adsorption ratio.
| Soil age (months) | Conductivity (dS m^-1^) | Chloride (mEq L^-1^) | Sodium (mEq L^-1^) | pH | Calcium+magnesium (mEq L^-1^) | Olsen P (mg kg^-1^) | SAR | Potassium (mg kg^-1^) | Iron (mg kg^-1^) | Zinc (mg kg^-1^) | Manganese (mg kg^-1^) | Boron (mg L^-1^) | Ammonium (mg kg^-1^) | Nitrate (mg kg^-1^) | UV 254 (nm) | TOC |
| --- | --- | --- | --- | --- | --- | --- | --- | --- | --- | --- | --- | --- | --- | --- | --- | --- |
| 0 | 1.91 | 13.24 | 9.25 | 7.7 | 10.80 | 25.88 | 3.98 | 54.0 | 12.60 | 2.80 | 21.10 | 0.081 | 3.60 | 11.178 | 0.603 | 14.13 |
| 0 | 1.64 | 11.44 | 7.25 | 7.7 | 9.68 | 44.60 | 3.30 | 133.8 | 11.48 | 2.74 | 18.60 | 0.140 | 9.90 | 18.906 | 0.557 | 16.73 |
| 0 | 1.09 | 5.68 | 1.90 | 7.6 | 7.68 | 35.84 | 0.75 | 351.6 | 11.22 | 3.00 | 24.10 | 0.162 | 8.50 | 11.350 | 0.591 | 17.06 |
| 6 | 1.29 | 8.00 | 6.37 | 7.6 | 7.28 | 27.56 | 3.34 | 51.4 | 10.50 | 2.60 | 17.50 | 0.126 | 7.10 | 8.015 | 0.641 | 14.84 |
| 6 | 1.17 | 6.32 | 5.50 | 7.7 | 6.64 | 39.56 | 3.02 | 107.7 | 12.30 | 2.90 | 20.20 | 0.111 | 8.10 | 10.660 | 0.811 | 19.49 |
| 6 | 1.37 | 8.28 | 6.00 | 7.7 | 8.20 | 42.52 | 2.96 | 136.9 | 10.20 | 2.70 | 17.80 | 0.140 | 7.80 | 7.141 | 0.500 | 13.30 |
| 18 | 1.46 | 9.52 | 6.87 | 7.6 | 8.28 | 23.00 | 3.38 | 56.7 | 13.00 | 2.82 | 17.90 | 0.105 | 5.00 | 12.178 | 0.566 | 16.43 |
| 18 | 1.46 | 8.68 | 7.12 | 7.7 | 8.50 | 36.92 | 3.46 | 192.1 | 14.50 | 3.50 | 22.90 | 0.123 | 7.80 | 18.676 | 0.551 | 16.54 |
| 18 | 1.05 | 5.68 | 1.95 | 7.6 | 7.22 | 29.64 | 0.84 | 231.8 | 11.00 | 2.50 | 18.20 | 0.110 | 12.00 | 8.751 | 0.629 | 16.72 |
| 30 | 1.3 | 7.68 | 6.50 | 7.6 | 6.98 | 32.00 | 3.48 | 106.2 | 12.90 | 2.60 | 19.00 | 0.113 | 7.00 | 13.834 | 0.902 | 21.34 |
| 30 | 1.53 | 10.4 | 7.37 | 7.6 | 8.64 | 25.28 | 3.55 | 46.9 | 12.50 | 2.50 | 19.00 | 0.109 | 7.10 | 18.135 | 0.544 | 17.01 |
| 30 | 1.60 | 9.88 | 8.12 | 7.6 | 8.86 | 29.88 | 3.86 | 86.1 | 15.80 | 4.00 | 22.00 | 0.125 | 9.90 | 19.918 | 0.589 | 15.63 |
| 42 | 1.45 | 9.44 | 7.12 | 7.5 | 8.06 | 27.28 | 3.55 | 40.4 | 13.10 | 2.74 | 15.80 | 0.106 | 7.10 | 6.566 | 0.699 | 20.99 |
| 42 | 1.36 | 8.76 | 6.62 | 7.5 | 7.36 | 26.60 | 3.45 | 56.7 | 14.40 | 3.00 | 20.50 | 0.110 | 7.80 | 16.399 | 0.577 | 18.39 |
| 42 | 1.49 | 9.24 | 6.87 | 7.4 | 8.74 | 37.64 | 3.29 | 75.3 | 15.50 | 4.00 | 26.40 | 0.109 | 6.40 | 13.604 | 1.022 | 22.12 |

### Slide 2
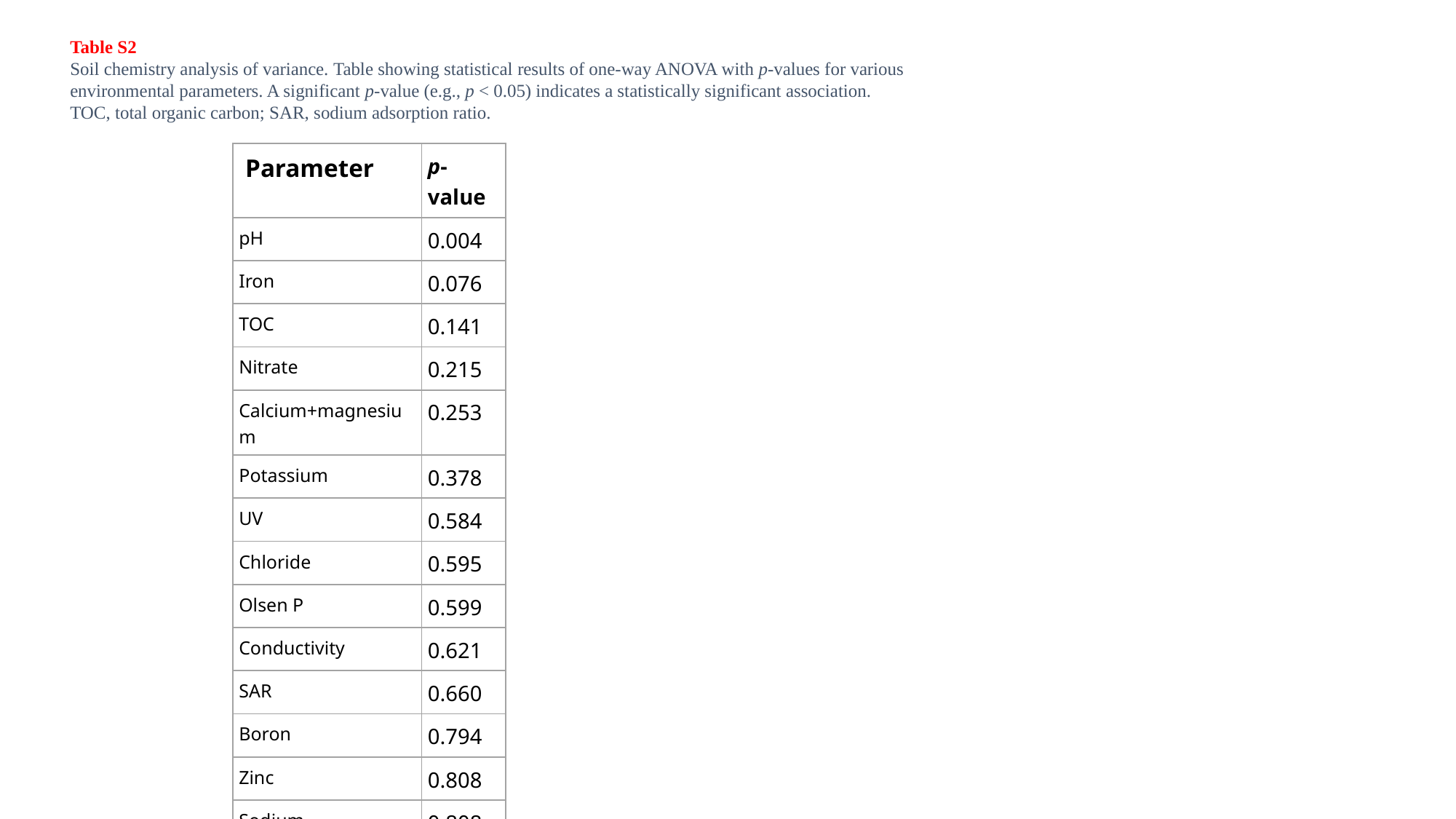

Table S2
Soil chemistry analysis of variance. Table showing statistical results of one-way ANOVA with p-values for various environmental parameters. A significant p-value (e.g., p < 0.05) indicates a statistically significant association. TOC, total organic carbon; SAR, sodium adsorption ratio.
| Parameter | p-value |
| --- | --- |
| pH | 0.004 |
| Iron | 0.076 |
| TOC | 0.141 |
| Nitrate | 0.215 |
| Calcium+magnesium | 0.253 |
| Potassium | 0.378 |
| UV | 0.584 |
| Chloride | 0.595 |
| Olsen P | 0.599 |
| Conductivity | 0.621 |
| SAR | 0.660 |
| Boron | 0.794 |
| Zinc | 0.808 |
| Sodium | 0.808 |
| Manganese | 0.830 |

### Slide 3
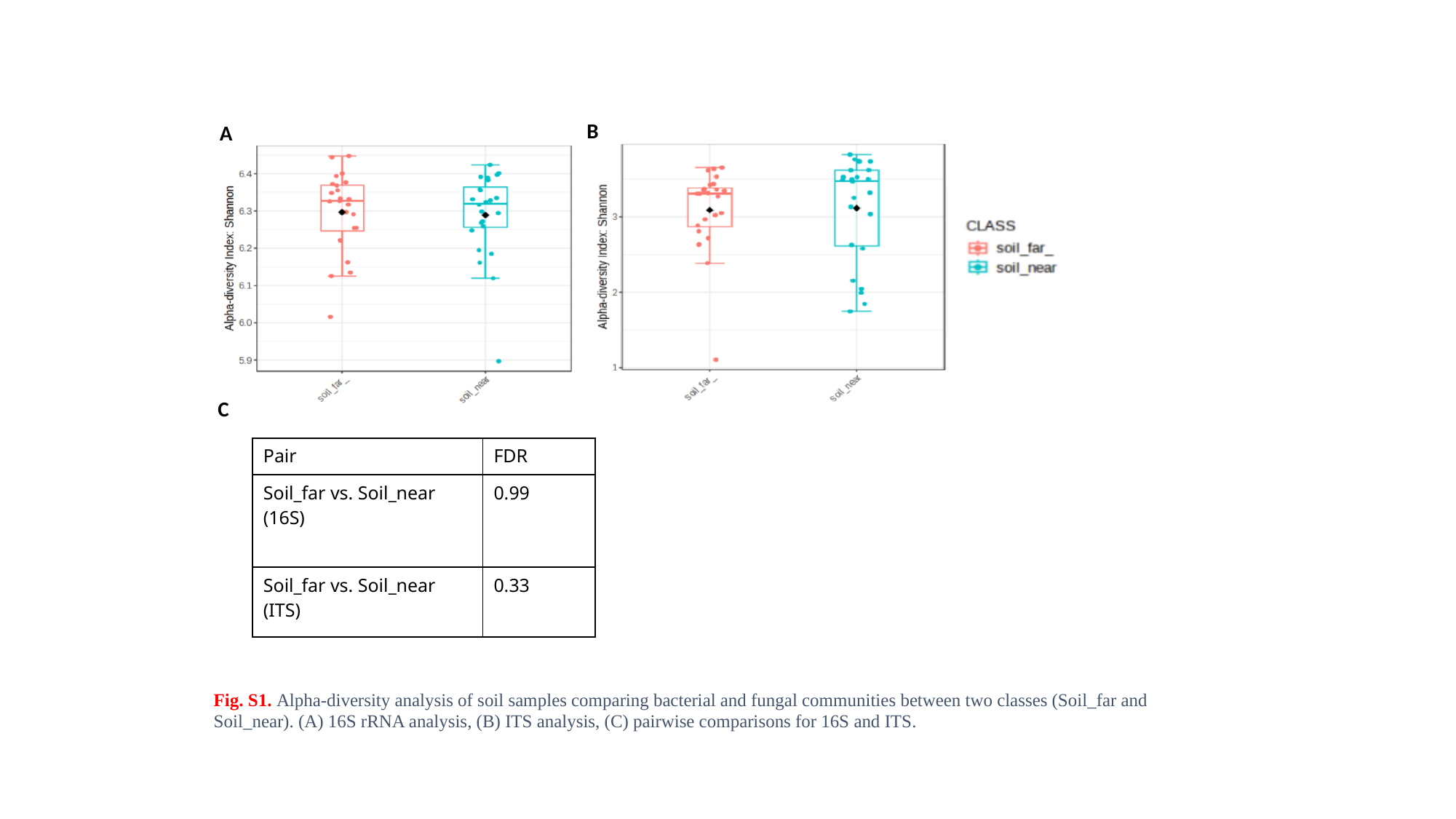

B
A
C
Fig. S1. Alpha-diversity analysis of soil samples comparing bacterial and fungal communities between two classes (Soil_far and Soil_near). (A) 16S rRNA analysis, (B) ITS analysis, (C) pairwise comparisons for 16S and ITS.
| Pair | FDR |
| --- | --- |
| Soil\_far vs. Soil\_near (16S) | 0.99 |
| Soil\_far vs. Soil\_near (ITS) | 0.33 |

### Slide 4
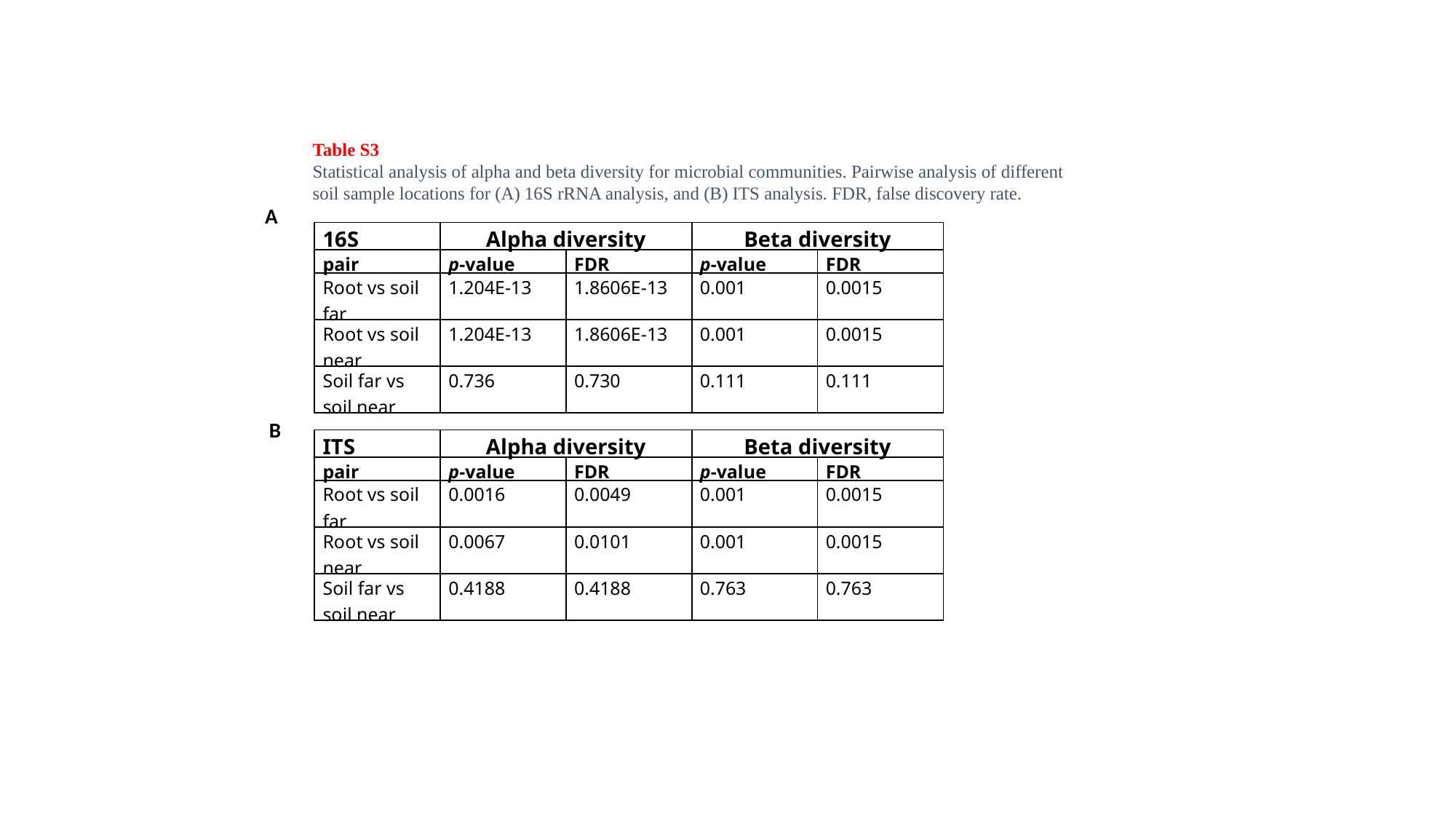

Table S3
Statistical analysis of alpha and beta diversity for microbial communities. Pairwise analysis of different soil sample locations for (A) 16S rRNA analysis, and (B) ITS analysis. FDR, false discovery rate.
A
| 16S | Alpha diversity | | Beta diversity | |
| --- | --- | --- | --- | --- |
| pair | p-value | FDR | p-value | FDR |
| Root vs soil far | 1.204E-13 | 1.8606E-13 | 0.001 | 0.0015 |
| Root vs soil near | 1.204E-13 | 1.8606E-13 | 0.001 | 0.0015 |
| Soil far vs soil near | 0.736 | 0.730 | 0.111 | 0.111 |
B
| ITS | Alpha diversity | | Beta diversity | |
| --- | --- | --- | --- | --- |
| pair | p-value | FDR | p-value | FDR |
| Root vs soil far | 0.0016 | 0.0049 | 0.001 | 0.0015 |
| Root vs soil near | 0.0067 | 0.0101 | 0.001 | 0.0015 |
| Soil far vs soil near | 0.4188 | 0.4188 | 0.763 | 0.763 |

### Slide 5
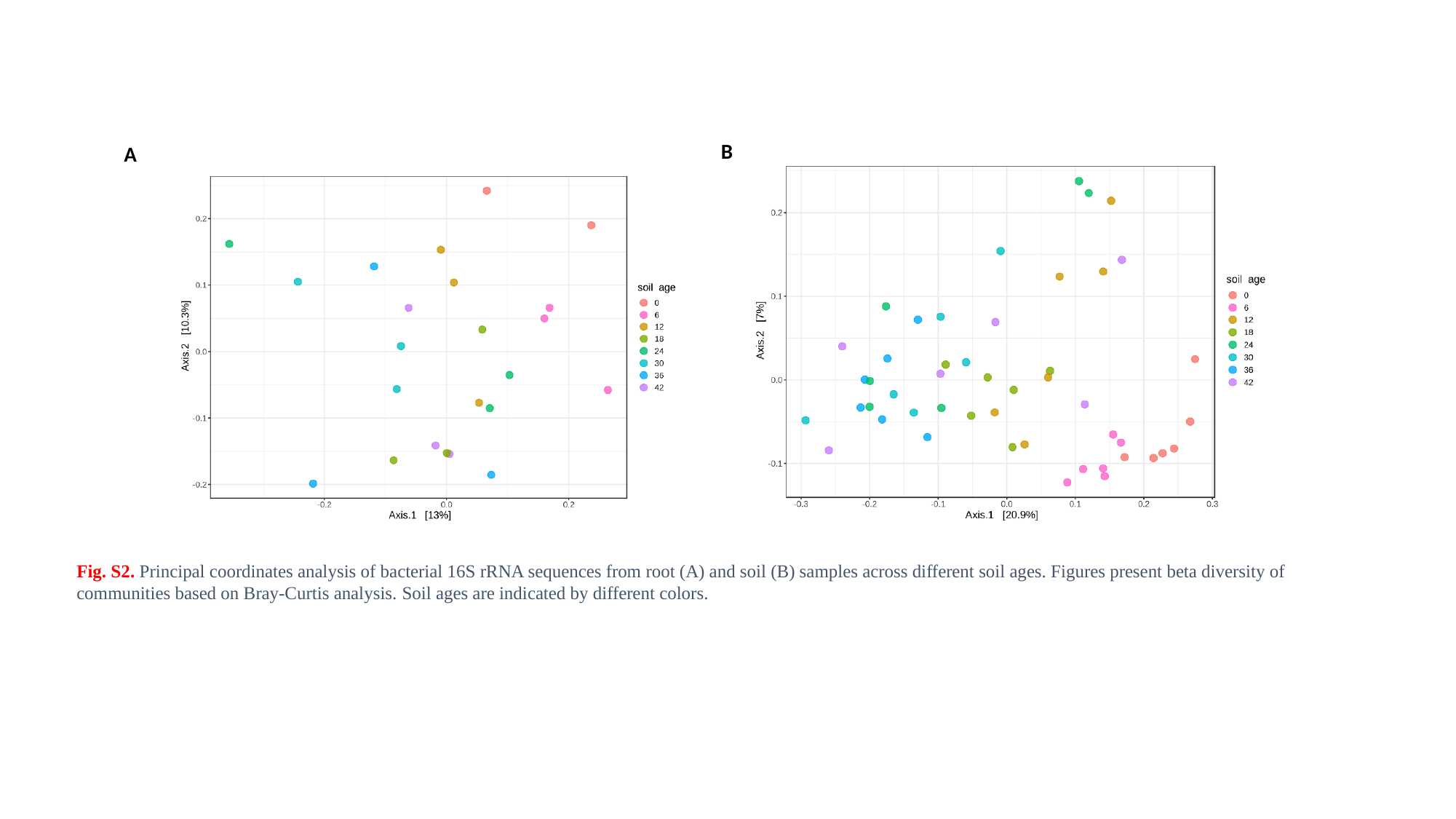

B
A
Fig. S2. Principal coordinates analysis of bacterial 16S rRNA sequences from root (A) and soil (B) samples across different soil ages. Figures present beta diversity of communities based on Bray-Curtis analysis. Soil ages are indicated by different colors.

### Slide 6
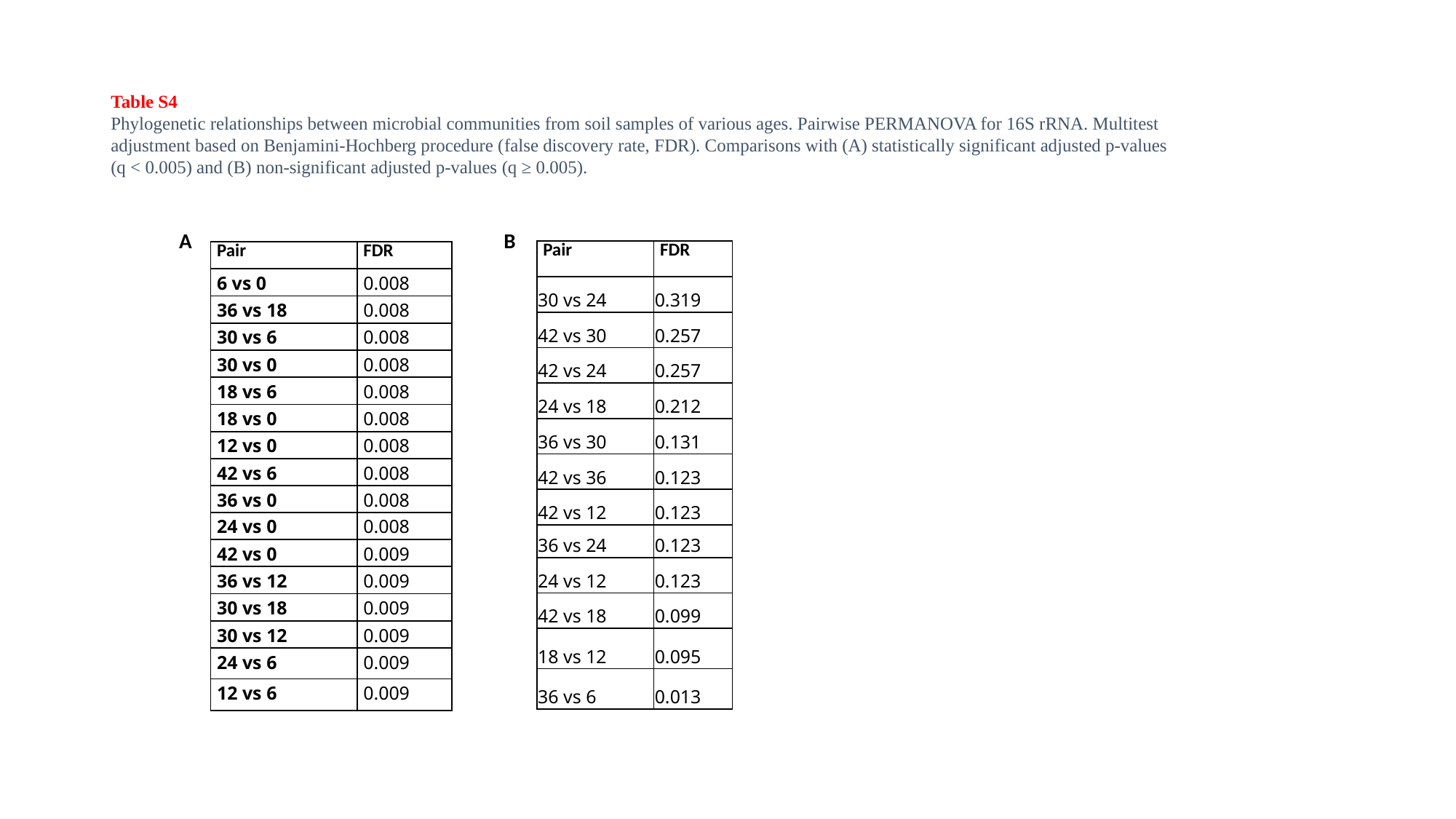

Table S4
Phylogenetic relationships between microbial communities from soil samples of various ages. Pairwise PERMANOVA for 16S rRNA. Multitest adjustment based on Benjamini-Hochberg procedure (false discovery rate, FDR). Comparisons with (A) statistically significant adjusted p-values (q < 0.005) and (B) non-significant adjusted p-values (q ≥ 0.005).
A
B
| Pair | FDR |
| --- | --- |
| 30 vs 24 | 0.319 |
| 42 vs 30 | 0.257 |
| 42 vs 24 | 0.257 |
| 24 vs 18 | 0.212 |
| 36 vs 30 | 0.131 |
| 42 vs 36 | 0.123 |
| 42 vs 12 | 0.123 |
| 36 vs 24 | 0.123 |
| 24 vs 12 | 0.123 |
| 42 vs 18 | 0.099 |
| 18 vs 12 | 0.095 |
| 36 vs 6 | 0.013 |
| Pair | FDR |
| --- | --- |
| 6 vs 0 | 0.008 |
| 36 vs 18 | 0.008 |
| 30 vs 6 | 0.008 |
| 30 vs 0 | 0.008 |
| 18 vs 6 | 0.008 |
| 18 vs 0 | 0.008 |
| 12 vs 0 | 0.008 |
| 42 vs 6 | 0.008 |
| 36 vs 0 | 0.008 |
| 24 vs 0 | 0.008 |
| 42 vs 0 | 0.009 |
| 36 vs 12 | 0.009 |
| 30 vs 18 | 0.009 |
| 30 vs 12 | 0.009 |
| 24 vs 6 | 0.009 |
| 12 vs 6 | 0.009 |

### Slide 7
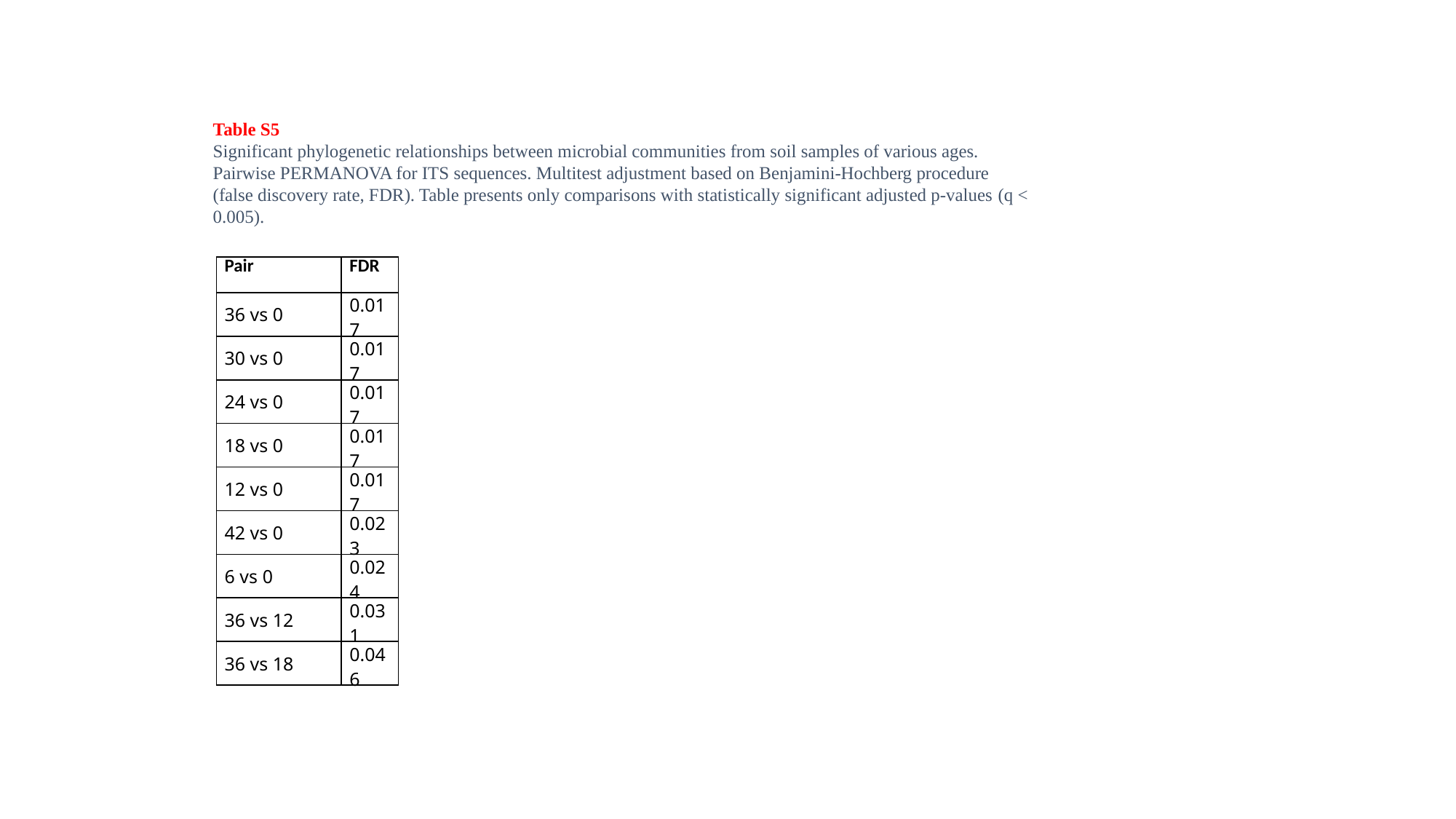

Table S5
Significant phylogenetic relationships between microbial communities from soil samples of various ages. Pairwise PERMANOVA for ITS sequences. Multitest adjustment based on Benjamini-Hochberg procedure (false discovery rate, FDR). Table presents only comparisons with statistically significant adjusted p-values (q < 0.005).
| Pair | FDR |
| --- | --- |
| 36 vs 0 | 0.017 |
| 30 vs 0 | 0.017 |
| 24 vs 0 | 0.017 |
| 18 vs 0 | 0.017 |
| 12 vs 0 | 0.017 |
| 42 vs 0 | 0.023 |
| 6 vs 0 | 0.024 |
| 36 vs 12 | 0.031 |
| 36 vs 18 | 0.046 |

### Slide 8
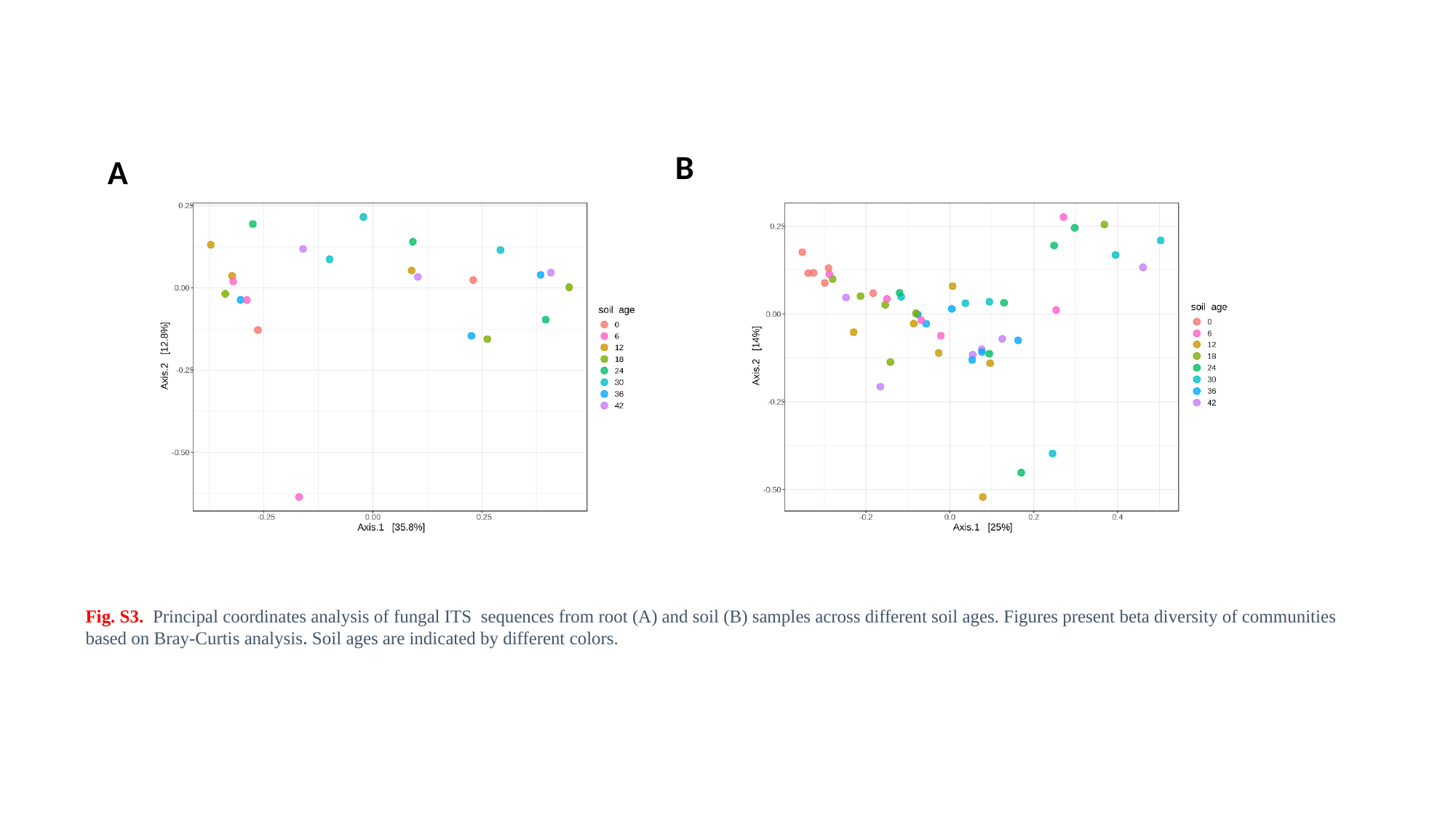

B
A
Fig. S3. Principal coordinates analysis of fungal ITS sequences from root (A) and soil (B) samples across different soil ages. Figures present beta diversity of communities based on Bray-Curtis analysis. Soil ages are indicated by different colors.

### Slide 9
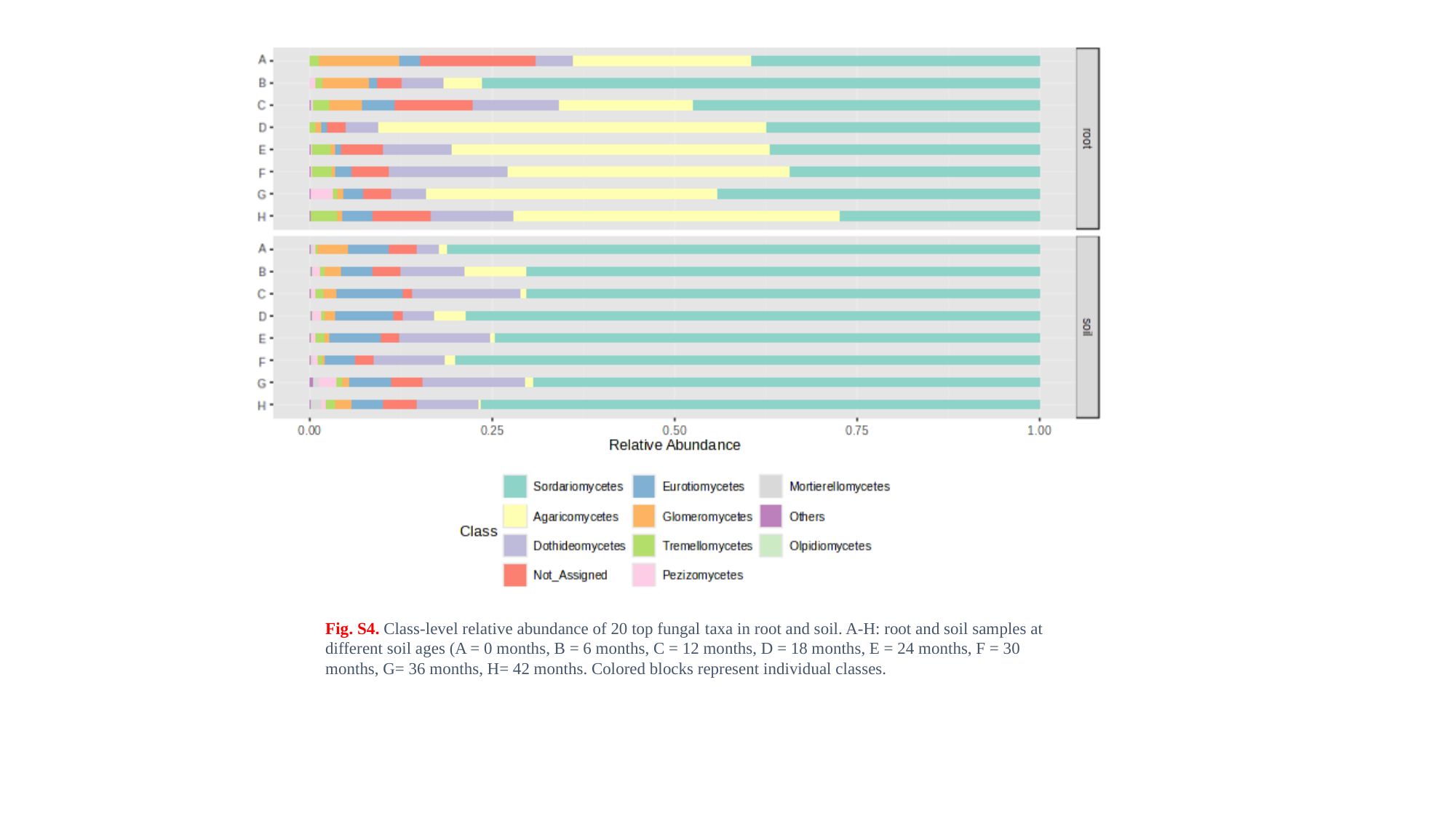

Fig. S4. Class-level relative abundance of 20 top fungal taxa in root and soil. A-H: root and soil samples at different soil ages (A = 0 months, B = 6 months, C = 12 months, D = 18 months, E = 24 months, F = 30 months, G= 36 months, H= 42 months. Colored blocks represent individual classes.
